# The unique *Brucella* effectors NyxA and NyxB target SENP3 to modulate the subcellular localisation of nucleolar proteins

**DOI:** 10.1101/2021.04.23.441069

**Authors:** Artur Louche, Amandine Blanco, Thais Lourdes Santos Lacerda, Claire Lionnet, Célia Bergé, Monica Rolando, Frédérique Lembo, Jean-Paul Borg, Carmen Buchrieser, Masami Nagahama, Jean-Pierre Gorvel, Virginie Gueguen-Chaignon, Laurent Terradot, Suzana P. Salcedo

## Abstract

The cell nucleus is a primary target for intracellular bacterial pathogens to counteract immune responses and hijack host signalling pathways to cause disease. The mechanisms controlling nuclear protein localisation in the context of stress responses induced upon bacterial infection are still poorly understood. Here we show that the *Brucella abortus* effectors NyxA and NyxB interfere with the host sentrin specific protease 3 (SENP3), which is essential for intracellular replication. Translocated Nyx effectors directly interact with SENP3 *via* a defined acidic patch identified from the crystal structure of NyxB, preventing its nucleolar localisation at the late stages of the infection. By sequestering SENP3, the Nyx effectors induce the cytoplasmic accumulation of the nucleolar AAA-ATPase NVL, the large subunit ribosomal protein L5 (RPL5) and the ribophagy receptor NUFIP1 in Nyx-enriched structures in the vicinity of replicating bacteria. This shuttling of ribosomal biogenesis-associated nucleolar proteins is negatively regulated by SENP3 and dependent on the autophagy-initiation protein Beclin1, indicative of a ribophagy-derived process induced during *Brucella* infection. Our results highlight a new nucleomodulatory function by two unique *Brucella* effectors, and reveal that SENP3 is a critical regulator of the subcellular localisation of multiple nucleolar proteins during *Brucella* infection, promoting intracellular replication.

## Introduction

The spatial organization of eukaryotic cells and accurate sub-cellular targeting of macromolecules is absolutely essential for the regulation of numerous cellular processes, namely cell growth, survival and stress responses. Indeed, mislocalisation of proteins in the cell has been associated with cellular stress and multiple diseases (1). The nucleus plays a critical role in sensing and orchestrating the overall host responses to stress. Several nuclear proteins have been shown to act as important stress-sensors in the cell, including the sentrin specific protease 3 (SENP3) which is also involved in regulation of cell cycle, survival pathways and ribosomal biogenesis (2). SENP3 functions as a processing enzyme for the SUMO2/3 precursor and as a deconjugase of SUMOylated substrates. SENP3 is mostly nucleolar and has been directly implicated in cellular adaptation to mild oxidative stress (3) and starvation (4). More recently, SENP3 was also proposed as a negative regulator of autophagy during nutritional stress (4). Autophagy, is one of the key stress-response mechanisms enabling cells to quickly adapt by engulfing cellular components such as damaged organelles (e.g. mitophagy and ER-phagy), toxic protein aggregates (proteophagy) or microbes (xenophagy) and directing them for degradation. Ribophagy is a selective type of autophagy involved in the degradation of ribosomes that disrupts ribosomal biogenesis and provides a swift source of nutrients for cells (5–7). Although multiple ribophagy pathways have been characterized depending on the specific stress signals involved, ribophagy was shown to be dependent on the autophagy-initiation protein Beclin1 (7). A single ribophagy receptor has been described to date (8), the nuclear fragile X mental retardation-interacting protein (NUFIP1), a nucleoplasmic protein first shown to be involved in 60S rRNA biogenesis in yeast (9). NUFIP1 is exported to the host cytoplasm in human cells upon starvation or mTOR inhibition, where it binds ribosomes and mediates their degradation (8). NUFIP1 is necessary for maintaining nucleoside/nucleotide levels and cell survival in the context of prolonged nutrient deprivation (8).

Bacterial pathogens can escape autophagy-mediated killing and, in some cases, modulate autophagy components and pathways to promote virulence. One such example is *Brucella abortus*, an intracellular pathogen that extensively replicates inside cells in a vacuole derived from the endoplasmic reticulum (ER), known as replicative *Brucella*-containing vacuole (rBCV). Although the role of autophagy in *Brucella* virulence remains poorly understood, several proteins have been implicated in different stages of the intracellular life cycle, notably the formation of rBCVs (Atg9 and WIPI-1) (10) and induction of autophagic BCVs mediating bacterial egress from infected cells (ULK and Beclin1) (11). The establishment of its replication niche requires the VirB/D type 4 secretion system (T4SS) that translocates effector proteins into host cells that specifically modulate cellular functions. Only very few *Brucella* effectors have been characterized to date, including some targeting innate immune responses (12–14) and the secretory pathway (15).

In this study we identify two new translocated effectors, NyxA and NyxB, that interact with the SENP3, leading to its subnuclear mislocalisation during infection. The crystal structure of NyxB and structure-based mutagenesis enabled us to pinpoint residues of the protein essential for SENP3 interaction. We show that NyxA and NyxB mediate the formation of *Brucella*-induced foci enriched in both effectors, the 60S ribosomal subunit protein RPL5, the AAA-ATPase NVL, and the ribophagy receptor NUFIP1. The formation of these structures is dependent on Beclin1 and is negatively regulated by SENP3, that *Brucella* sequesters to sustain intracellular replication. The modulation of nuclear functions during infection is an important virulence strategy shared by a growing number of bacterial pathogens (16). In most cases, this nuclear targeting results in a fine regulation of host gene expression or control of cell cycle to benefit host colonization and persistence. In this study we highlight a new nucleomodulation mechanism that promotes perturbation of the subcellular localisation of nucleolar proteins during bacterial infection.

## Results

### The newly identified *B. abortus* effectors NyxA and NyxB accumulate in cytoplasmic and nuclear structures

Bacterial effectors often contain eukaryotic-like domains to enable efficient modulation of cellular pathways. Several bacterial effectors rely on a carboxyl-terminal CAAX tetrapeptide motif (C corresponds to cysteine, A to aliphatic amino acids and X to any amino acid) as a lipidation site to facilitate membrane attachment (17–20), with the presence of at least one carboxy-terminal cysteine being the essential feature and the remaining motif more flexible (21). Previous work listed a subgroup of *Brucella* candidate effectors containing this kind of potential lipidation motif (19). We have recently confirmed that one of these *Brucella* proteins, BspL, is translocated into host cells during infection (22). Therefore, we set out to determine if two other *B. abortus* proteins, encoded by BAB1_0296 (BAB_RS17335) and BAB1_0466 (BAB_RS18145), could be translocated into host cells during infection. We relied on the TEM1 ß-lactamase reporter, widely used to assess translocation of *Brucella* effectors in RAW macrophage-like cells, allowing very high rates of infection. At 24h post-infection we observed that BAB1_0296 was efficiently translocated into host cells, in contrast to BAB1_0466 (Figure 1A and B). Hence, we can conclude that BAB1_0296 is likely to be a *B. abortus* effector, and we have named it as NyxA, inspired by Greek mythology. Nyx is the Greek personification of the night, daughter of Chaos, that seemed appropriate to us after remaining in the dark for so long regarding its function. It also alludes to its function as a Nucleomodulin ribophagy-exploiting protein. Genome analysis identified another gene, BAB1_1101 (BAB_RS21200), encoding for a protein with 82% identity to NyxA, but without a carboxy-terminal CAAX motif (Supplementary Figure 1A). For consistency, we have named it NyxB and found that it was also translocated by *B. abortus* into host cells, albeit to lower levels than NyxA at both 4 and 24h post-infection (Figure 1A and B and Supplementary Figure 1B). Both NyxA and NyxB are well conserved within the *Brucella* species and show little homology to other bacterial proteins. Paralogs of NyxA and B are present in members of the closely related genus *Ochrobactrum* from the *Brucellaceae* family.

**Fig. 1.**
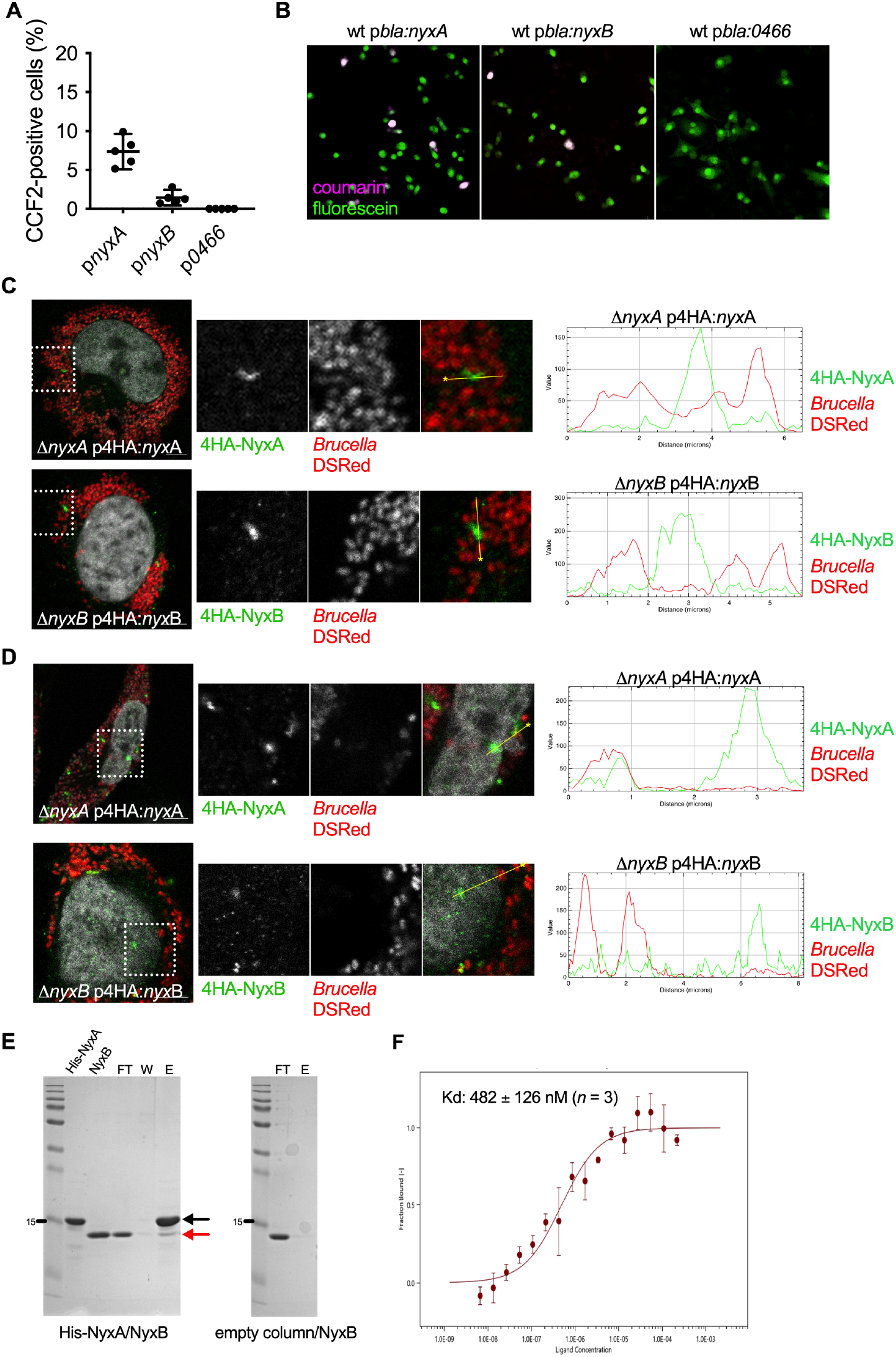
The *Brucella* NyxA and NyxB proteins are translocated into host cells during infection, accumulating in punctate or filament-like cytoplasmic and nuclear structures and can interact with each other. **(A)** RAW macrophage-like cells were infected for 24h with either *B. abortus* wild-type expressing TEM1 (encoded by the *bla* gene) fused with NyxA, NyxB or BAB1_0466. The percentage of cells with coumarin emission, which is indicative of translocation, was quantified after incubation with the CCF2-AM substrate. Data represent means ± 95% confidence intervals from 5 independent experiments, with more than 500 cells counted for each condition. Representative images for *B. abortus* wild-type carrying p*bla*:*nyxA*, p*bla*:*nyxB* or p*bla*:BAB1_0466 to exemplify the presence of effector translocation visible in coumarin-positive cells (violet) or absence of translocation. **(C)** Accumulation of 4HA-tagged NyxA (top) or 4HA-tagged NyxB (bottom) in cytoplasmic punctate or filament-like structures in HeLa cells infected for 48h with Δ*nyxA* or Δ*nyxB* strains expressing DSRed and the corresponding 4HA-tagged effector. The cell nucleus is visible with DAPI. A fluorescence intensity profile along a defined straight line across the 4HA-positive structures is included for each image, with the HA signal represented in green and bacterial signal in red. **(D)** Representative confocal microscopy images showing accumulation of 4HA-tagged NyxA (top) or 4HA-tagged NyxB (bottom) in nuclear punctate structures in HeLa cells infected for 48h with Δ*nyxA* or Δ*nyxB* strains expressing DSRed and the corresponding 4HA-tagged effector. **(E)** Pulldown experiment using purified NyxB against His-NyxA immobilised on a Ni NTA resin. An empty column was used as a control for non-specific binding. Interactions were visualised with coomassie blue stained gels. The flowthrough (FT), wash (W) and elution (E) fractions are shown for each sample and the molecular weights indicated (kDa). Eluted His-NyxA and NyxB are indicated with black and red arrows, respectively. **(F)** Microscale thermophoresis measuring the fraction of 20 nM of purified NyxA labelled with kit protein labelling RED-NHS binding to increasing concentrations of NyxB (6.67 nM to 219 μM). Data correspond to means ± standard deviations of 3 independent experiment. The obtained Kd is indicated.

To gain insight into the mechanism of translocation of NyxA and NyxB, we infected cells for 4h, a time point that enables a comparison between the wild-type and the Δ*virB9* mutant strain lacking a functional T4SS. NyxA translocation was significantly reduced in a *virB* mutant strain, suggesting dependency on the T4SS. We should note that a small number of cells infected with the Δ*virB9* expressing NyxA were positive for CCF2 compared to BAB1_0466, for which no translocation was detected suggesting a proportion of NyxA could also be translocated independently of the T4SS (Supplementary Figure 1B and C). Curiously, the translocation of NyxB was independent of the T4SS (Supplementary Figure 1B).

To confirm that NyxA was indeed translocated across the vacuolar membrane during infection, we constructed new strains with NyxA fused on its N-terminus with a 4HA epitope tag, successfully used for imaging bacterial effectors (23). We then infected HeLa cells, a well-characterised model of *B. abortus* infection nicely suited for microscopy studies. We could observe translocated 4HA-NyxA at 48h post-infection, accumulating in cytoplasmic structures in the vicinity of multiplying bacteria (Figure 1C). This was also the case for 4HA-NyxB (Figure 1C, bottom panel). Analysis of fluorescence intensity profiles along a defined straight line across the 4HA-positive structures confirmed the majority of the 4HA signal detected does not correspond to intra-vacuolar NyxA or NyxB. For both effectors, punctate and filament-like structures were observed (Supplementary Figure 1D) and as early as 24h post-infection. Equivalent NyxA structures were observed with an N-terminal 3Flag (Supplementary Figure 1E), suggesting these are not an artefact due to the 4HA tag. The 4HA-NyxA and NyxB positive structures could also be detected in the nucleus at 48 and 65h post-infection (Figure 1D), suggesting these effectors may also be targeting the host nuclei, particularly at late stages of the infection.

### NyxA and NyxB interact with each other and target the same cellular compartments

Imaging of translocated NyxA and NyxB suggested that these effectors may have an identical subcellular localisation. To investigate this possibility, we ectopically expressed NyxA and NyxB with different tags. We found that HA-tagged NyxA and NyxB predominantly accumulated in nuclear aggregates (Supplementary Figure 2A, and B), with a few cytoplasmic vesicular structures visible in most cells. Interestingly, 4HA-tagged NyxA and NyxB were mostly found in the cytoplasm in structures resembling what we observed during infection (Supplementary Figure 2A, and B). This suggests that the presence of the 4HA reduces the nuclear import of NyxA and NyxB, revealing its initial cytoplasmic location. This hypothesis was not confirmed in infected cells as effectors tagged with a single HA epitope were not detectable at any of the time-points analysed.

The co-transfection of HA-NyxA and myc-NyxB showed substantial co-localisation levels, suggesting that these two proteins target the same cellular compartments (Supplementary Figure 2C). To determine if these two effectors could interact, we purified them both and carried out pull-down experiments *in vitro*. We found that His-V5-NyxA could pull-down purified untagged NyxB (Figure 1E) and vice-versa (Supplementary Figure 3A and B). Furthermore, microscale thermophoresis confirmed this interaction and determined a Kd of 482 ± 126 nM (Figure 1F). Together, these results show NyxA and NyxB interact and can accumulate in cytoplasmic structures as well as in the nucleus.

### The Nyx effectors interact with SENP3, which is necessary for efficient *B. abortus* intracellular multiplication

To investigate the potential role of NyxA and NyxB during infection, we constructed deletion mutants in *B. abortus* 2308. The overall intracellular bacterial counts of the single or double mutant strains were equivalent to the wild-type, suggesting that the lack of these effectors does not significantly impact establishment of the replicative niche (Supplementary Figure 4), often observed for *Brucella* and other bacterial pathogens as *Legionella* with high effector redundancy. Therefore, to characterize the function of NyxA and NyxB we identified their potential host-interacting partners, by performing a yeast two-hybrid screen of NyxA against a library of human proteins (Supplementary Table 1). One of the main proteins identified was SENP3, and in view of its nuclear localisation and its diverse range of functions in key cellular processes and response to stress, we focused on this potential target.

SENP3 belongs to a family of cysteine proteases that share a conserved catalytic domain, characterised by a papain-like fold (24). The variable N-terminal region often contributes to intracellular targeting of the protease. In the case of SENP3, the N-terminal region is implicated in nucleolar targeting as deletion of the residues 76 to 159 prevents nucleolar accumulation (25). This region encodes two basic amino acid stretches predicted as nucleolar localisation sequences, overlapping with the NPM1 binding domain and an mTOR phosphorylation site. The phosphorylation of SENP3 by the mTOR kinase facilitates interaction with NPM1, which enables subsequent nucleolar shuttling (25).

To confirm the interaction between SENP3 and the Nyx effectors, we attempted to purify SENP3. We were not successful and instead focused on the purification of the N-terminal region of SENP3 that encompassed all the yeast two-hybrid hits (SENP_7-159_). Both His-V5-tagged NyxA and NyxB were able to pull-down SENP_7-159_, confirming a direct interaction of these effectors with the N-terminus domain of SENP3 (Figure 2A). No unspecific binding to the column was detected (right panel, Figure 2A). We cannot exclude the involvement of other regions of SENP3, but our data show SENP_7-159_ is sufficient for this interaction.

**Fig. 2.**
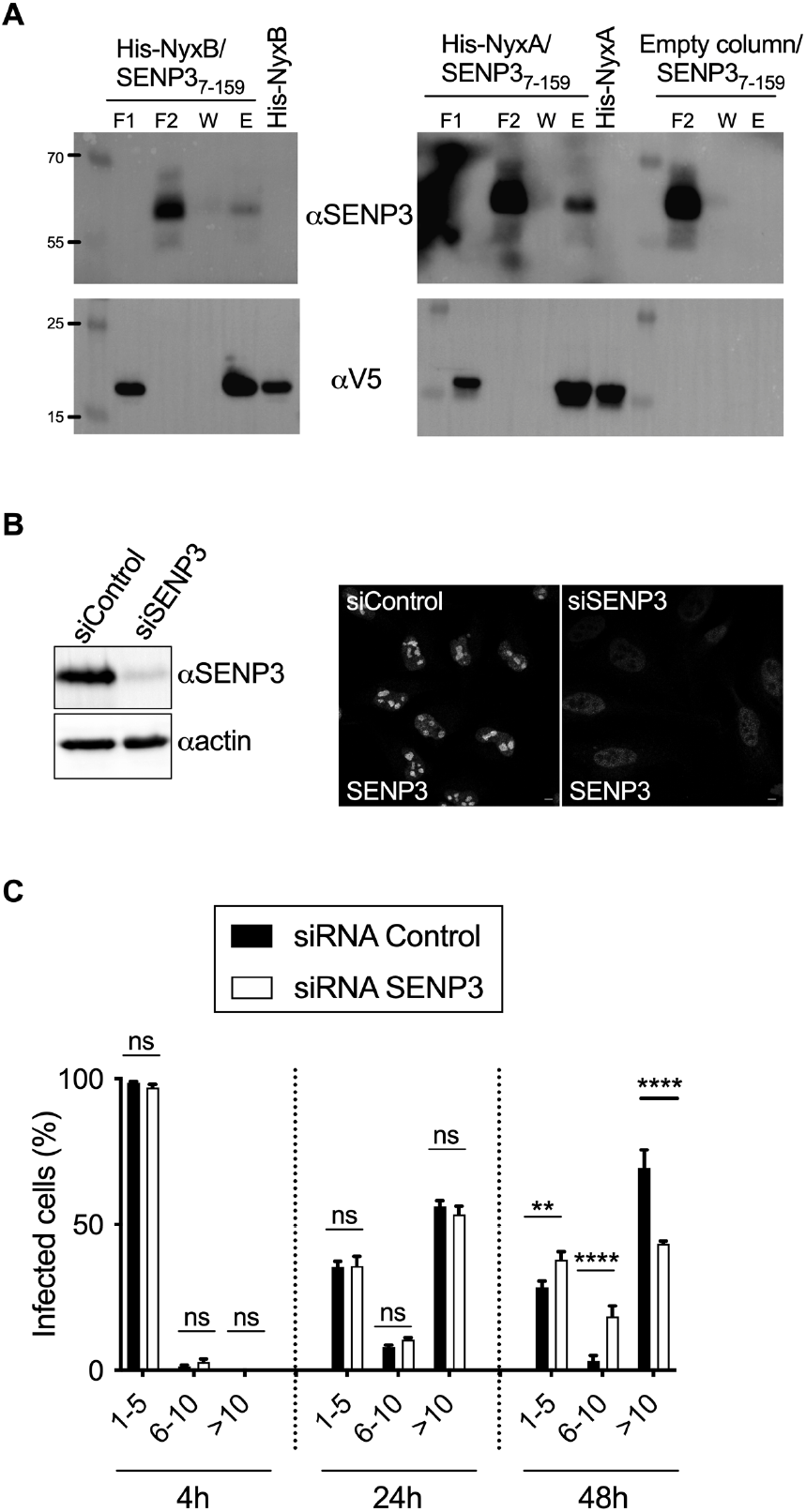
The Nyx effectors interact with host protease SENP3, essential for efficient *B. abortus* intracellular multiplication. **(A)** Pull-down assay with the N-terminal region of SENP3 from amino acid 7 until 159 (SENP_7-159_) against His-V5-NyxA or His-V5-NyxB immobilised on Ni NTA resins. An empty column was used as a control for non-specific binding and purified His-NyxA and His-Nyx-B inputs are shown. Interactions were visualised by western blotting using anti-SENP3 antibody, and column binding with anti-V5 (lower blot). Non-bound fractions (F1 and F2), last wash (W) and elution (E) are shown for each sample and the molecular weights indicated (kDa). **(B)** Western blot of HeLa cell lysate treated with siRNA control (siControl) or siRNA SENP3 (siSENP3) for 72h. Membrane was probed with an anti-SENP3 antibody followed by anti-actin for loading control. Depletion was also verified by microscopy, showing a predominant nucleolar localisation of SENP3 in control cells which is strongly reduced in siSENP3 treated cells. Scale bar is 5 μm. HeLa cells depleted for SENP3 or treated with the control siRNA were infected with wild-type *B. abortus* expressing DSRed and the percentage of cells with either 1 to 5, 6 to 10 or more than 10 bacteria per cell was quantified by microscopy at 2, 24 or 48h post-infection. Data correspond to means ± 95% confidence intervals from 3 independent experiments, with more than 500 cells being counted for each siRNA treatment at each time-point. A two-way ANOVA with Bonferroni correction was used to compare the bacterial counts obtained in siControl treated cells with siSENP3 depleted cells, for each subgroup (1-5, 6-10 or >10 bacteria/cell) at each time-point.

As SENP3 seems to be the host target of both Nyx effectors, we determined its relevance during infection. The depletion of SENP3 was efficiently achieved after treatment with siRNA (Figure 2B), after which cells were infected with wild-type *B. abortus*. We observed a reduction in the percentage of cells with more than ten bacteria at 48h post-infection when SENP3 was depleted but not at earlier stages (Figure 2C). Therefore, in the late stages of the infection, SENP3 was required for *B. abortus* to multiply efficiently inside cells.

### The NyxB structure defines a novel family of effectors

To gain further insight into the function of NyxA and NyxB and their interaction with SENP3, we solved the crystal structure of NyxB at 2.5Å (Figure 3A, Supplementary Table 2). NyxA did not form crystals in any of the conditions tested. The asymmetric unit of the crystal contains twelve monomers of NyxB but no significant differences were found between them and thus only the structure of subunit A is described hereafter. The NyxB model encompasses residues 17 to 134 suggesting that residues 1 to 16 are flexible. NyxB has a mixed α-β fold with five β strands and six α-helices with a core made of helices α2 to α4 and a small curved β-sheet formed by β3, β4 and β5. The longest helix α4 interacts with α2 and α6 and is connected to the core via a loop containing two short 3_10_ helices designated α3a and α3b (Figure 3A). Helix α1 is loosely associated with the rest of the protein core and is positioned by the preceding and following loops. In particular, a β-hairpin formed by β1 and β2 packs against α5 and anchors α1 to the protein core. Search for structural homologues (DALI server and EMBL fold (26)) did not reveal any significant homology and thus makes of NyxB structure a prototype for this protein family. However, the C-terminal part of the protein showed some similarity with a number of nucleic acid binding proteins, including Daschung, Ski or ribosomal protein RPL25 containing a winged helix domain. These similarities all match NyxB helices α2, α4 and α6 with the winged motif formed by strands β4 and β5.

**Fig. 3.**
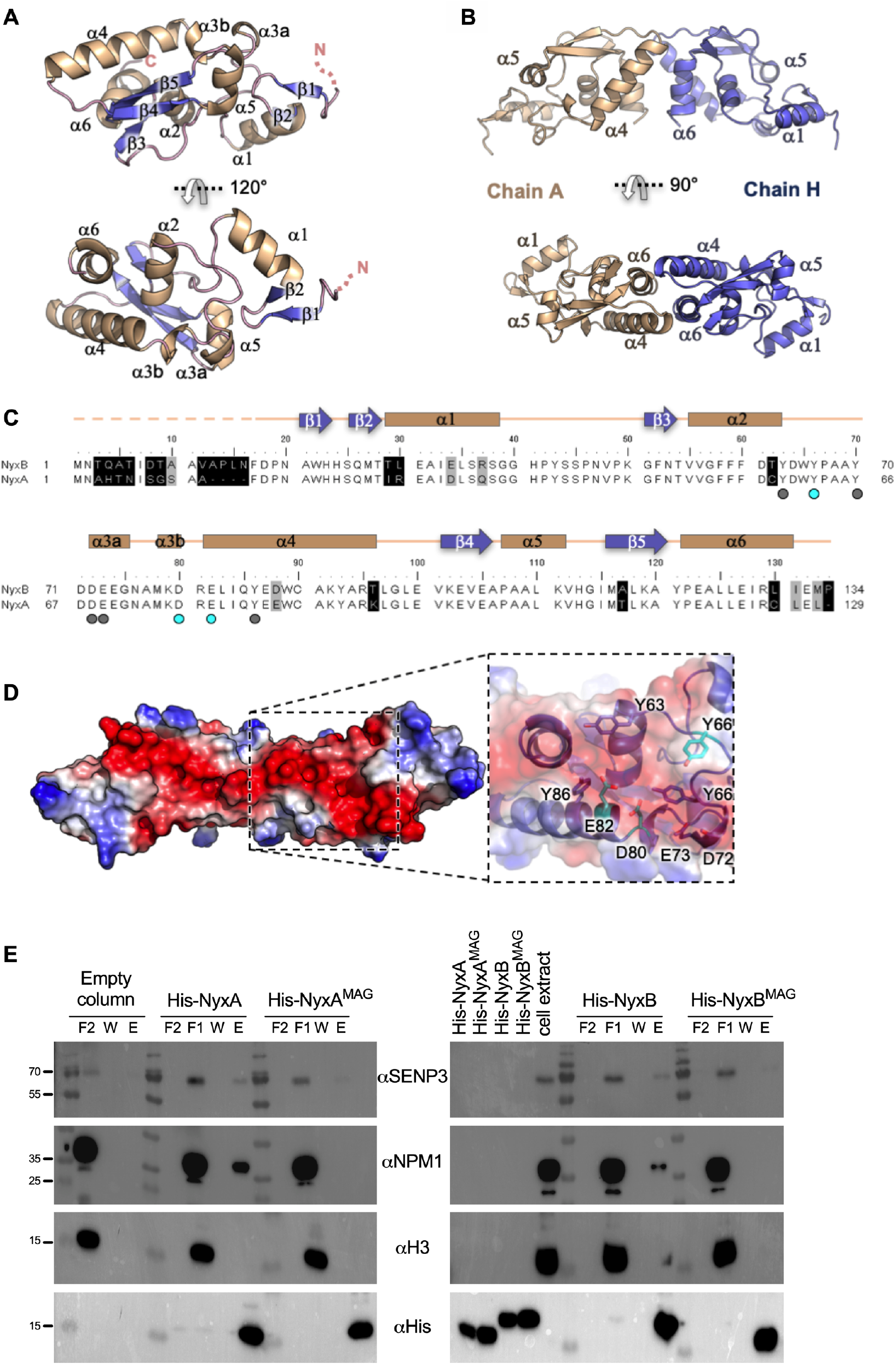
The NyxB structure defines a novel family of effectors and allowed identification of the SENP3 interacting groove. **(A)** Two views of the NyxB monomer depicted in ribbon with helices coloured in wheat, strands in blue and loops in pink. **(B)** Two views of the NyxB dimer. **(C)** Structure-based sequence alignment of NyxB and NyxA. Secondary structure elements are indicated above the sequences. Identical residues are not shaded, residues shaded in black and grey are non-conserved and conserved, respectively. Dots indicate residues identified in the acidic patch and cyan dots indicate acidic groove mutants (MAG). **(D)** Surface representation of NyxB dimer coloured according to electrostatic potential (red negative, blue positive) showing the extended acidic patch. The inset shows a close-up view of the area with residues’ side chains displayed as ball-and-sticks and mutated residues (E82,Y66 and D80) coloured in cyan. **(E)** Pull-down assay with His-NyxA, His-NyxB or the specific catalytic mutants (His-NyxA^MAG^ or His-NyxB^MAG^) immobilised on Ni NTA resins that were incubated with a HeLa cell extract. Empty column was used as a control for non-specific binding. Interactions with endogenous SENP3, NPM1 or Histone 3 (H3) were visualised by western blotting using the corresponding antibody, and column binding with anti-His (lower blot). Non-bound fractions (F1 and F2), last wash (W) and elution (E) are shown for each sample and the molecular weights indicated (kDa). The cell extract and the different purified Nyx inputs are also shown.

Size exclusion chromatography coupled to multi-angle light scattering (SEC-MALS) experiments indicate that both NyxA and NyxB form dimers (Supplementary Figure 5A). Two putative dimers (dimer 1 and dimer 2) were identified in the asymmetric unit (Supplementary Figure 5B). The association of dimer 1 (chain A and H) buries a total of 530 Å^2^ (Figure 3B and Supplementary Figure 5B) while dimer 2 (chain A and J) relies on fewer interactions burying a total 400 Å^2^ (Supplementary Figure 5B). Small angle X-ray scattering on NyxB and NyxA proteins (Supplementary data, Supplementary Table 3 and Supplementary Figure 5C and D) clearly showed that the two proteins adopt predominantly dimer 1 conformation in solution. This assembly relies on reciprocal hydrophobic and electrostatic interactions between α4 of one subunit and α6 of the other subunit and between the two α4-β4 loops (Figure 3A). These results show that NyxA and NyxB are members of the same family and share a number of properties, including their tertiary and quaternary structures.

### Identification of the Nyx-SENP3 interacting residues

Taking advantage of the structural information of NyxB, we searched for potential interaction sites with SENP3. Analysis of the NyxB surface revealed an acidic pocket delineated by residues Y66, D80 and E82 within an acidic patch consisting of amino acids D72, E73, Y70, Y86 and Y63, residues that are strictly conserved in NyxA (Figure 3C). In the context of the dimer, these surfaces are juxtaposed to form an extended concave negatively charged area of around 2000 Å^2^ (Figure 3D). To significantly modify the charge of the protein we generated a triple mutant Y66R, D80R and E82R to obtain His-NyxB^MAG^, where “MAG” stands for mutated acidic groove and Y62R, D76R and E78R to obtain His-NyxA^MAG^. Purified NyxA or NyxB were able to pull-down endogenous SENP3 from a HeLa cell extract confirming their interactions (Figure 3E). A decreased ability for both His-NyxA^MAG^ and His-NyxB^MAG^ to interact with SENP3 was observed (Figure 3E). However, we could only detect a small amount of endogenous SENP3 in the cell extract with our antibody. As it is well established that when SENP3 is pulled-down from a cell extract, its major cellular partner NPM1 (27) can easily be detected by western blotting, we next probed the same membrane with an antibody against NPM1. NPM1 was not pulled-down by His-NyxA^MAG^ and His-NyxB^MAG^, confirming that this mutation strongly impaired their ability to bind the complex SENP3-NPM1 (Figure 3E). As a negative control, the membrane was also probed for Histone 3, an abundant nuclear protein that did not bind to neither NyxA nor NyxB (Figure 3E). Together, these results confirm SENP3 as a target of the *Brucella* Nyx effectors and identify the acidic groove responsible for this interaction *in vitro*.

### The *Brucella* Nyx effectors induce delocalisation of SENP3

After showing that NyxA and NyxB interact directly with SENP3, a eukaryotic protein mainly found in the nucleoli (Figure 4A, top panel), we were intrigued by what impact these effectors could have on SENP3. Ectopic expression of either HA-tagged NyxA or NyxB resulted in a marked reduction of endogenous nucleolar SENP3, which instead formed aggregates in the nucleoplasm (Figure 4A and B). As SENP3 redistribution from nucleoli to the nucleoplasm could be due to starvation or mild oxidative stress (25,28), we ectopically expressed the mutant HA-NyxA^MAG^, unable to interact with SENP3. The mutation of the acidic interaction groove impaired delocalisation of SENP3 by NyxA and to a lesser extent NyxB, confirming this effect was due to direct interaction with SENP3. Consistently, analysis of the same images for co-localisation of SENP3 with HA-tagged Nyx proteins revealed important recruitment, dependent on the acidic groove (Figure 4C).

**Fig. 4.**
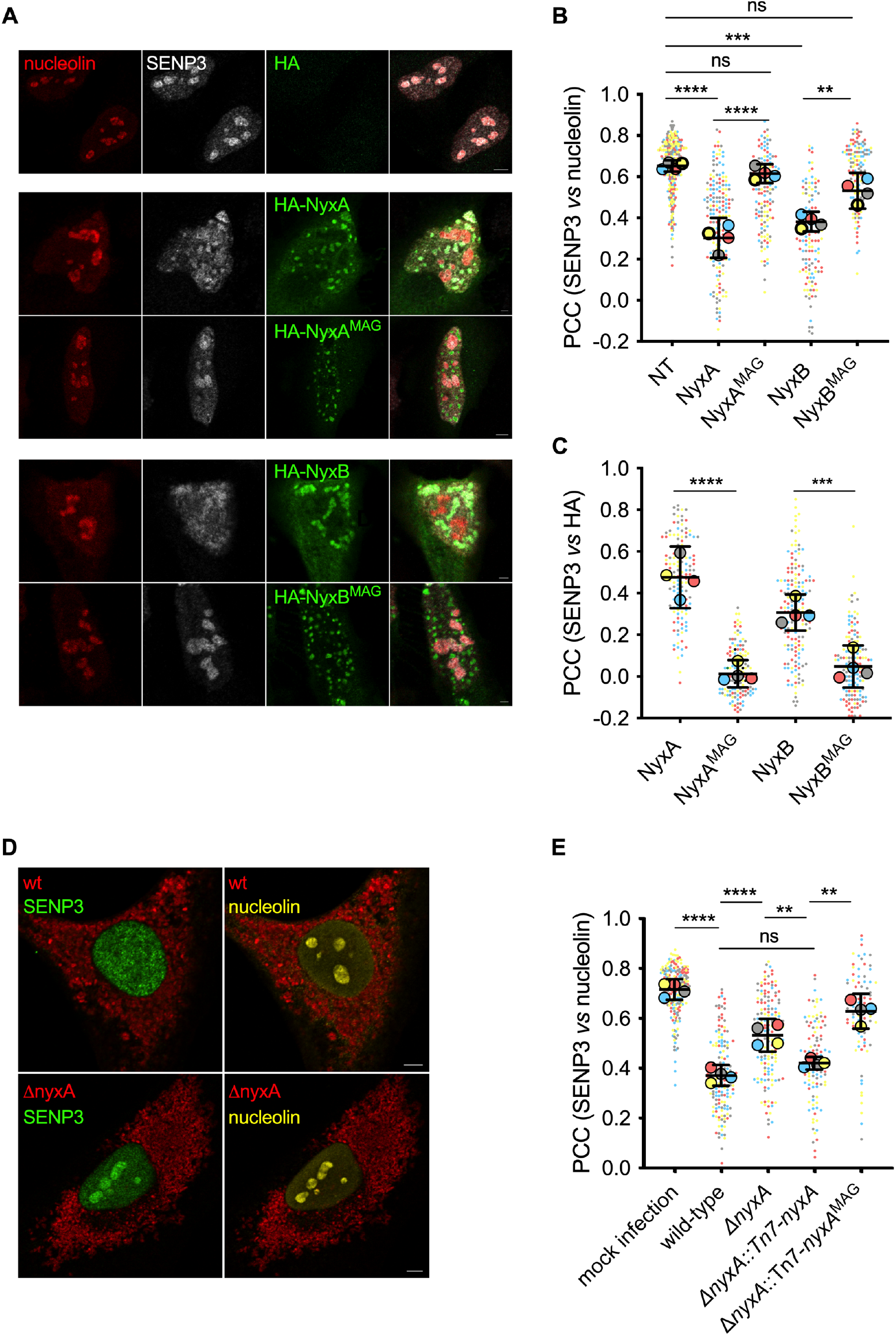
The *Brucella* Nyx effectors directly perturb the SENP3 nucleolar localisation in host cells, including during infection. **(A)** Representative confocal microscopy images of HeLa cells expressing the HA empty vector, HA-NyxA, HA-NyxA^MAG^, HA-NyxB and HA-NyxB^MAG^. Nucleolin (red), SENP3 (white) and HA (green) were revealed with specific antibodies. **(B)** Quantification of the Pearson correlation coefficient of SENP3 versus nucleolin (see methods for plugin description). Data are represented as means ± 95% confidence intervals from 4 independent experiments. Each experiment is colour coded and all events counted are shown. Data were analysed using one way ANOVA by including all comparisons with Tukey’s correction. Not all comparisons are shown. All the cells quantified are shown in the format of SuperPlots, with each colour representing an independent experiment and its corresponding mean (N = 4). **(C)** The same data set was used for quantification of the Pearson correlation coefficient of SENP3 versus HA to assess recruitment. **(D)** Representative confocal microscopy images of HeLa cells infected for 48h with wild-type DSRed expressing *B. abortus* or Δ*nyxA*, with nucleolin (yellow) and SENP3 (green). **(E)** Quantification of the Pearson’s coefficient of SENP3 versus nucleolin in HeLa cells infected for 48h with either *B. abortus* wild-type or Δ*nyxA*, its complemented strain Δ*nyxA*::Tn7-*nyxA* or a complementing strain expressing the mutated acidic groove responsible for interaction with SENP3 (Δ*nyxA*::Tn7-*nyxA*^MAG^). Data are represented and were analysed as in (B). Not all comparisons are shown. All microscopy images displayed have scale bars corresponding to 5 μm.

Next, we investigated the prevalence of these phenotypes during infection. We infected HeLa cells with either the wild-type *B. abortus* strain, the mutant lacking *nyxA*, or a complemented strain expressing *nyxA* from the chromosome under the control of its promoter.

Analysis of the level of SENP3 retained in the nucleoli during infection showed a significant lack of nucleolar localisation of SENP3 in cells infected with the wild-type *B. abortus* strain in contrast with the Δ*nyxA* strain (Figure 4D and E). The wild-type phenotype was partially restored with the complemented strain but not with a Δ*nyxA* strain expressing *nyxA*^MAG^. Lack of complementation observed for the strains expressing NyxA^MAG^ was not due to lack of translocation, as TEM-NyxA^MAG^ and TEM-NyxB^MAG^ were both efficiently translocated during infection (Supplementary Figure 6A). This result shows that NyxA’s interaction with SENP3 during infection prevents its accumulation in the nucleoli. In the case of NyxB, we could also observe a statistically significant increase of SENP3 in the nucleoli in cells infected with the Δ*nyxB* strain compared to cells infected with wild-type *B. abortus*, although to a lesser extent than what we observed for Δ*nyxA*. However, we could not fully complement this phenotype (Supplementary Figure 6B), possibly due to the low sensitivity of this microscopy approach combined with a weaker phenotype. As expected, a strain lacking both genes encoding for NyxA and NyxB could not mislocalise SENP3 as the wild-type strain (Supplementary Figure 6B). Representative images of all strains are shown in the Supplementary Figure 6C.

### Nyx effectors induce cytoplasmic accumulation of the nucleolar protein NVL in *Brucella*-induced foci (Bif)

One of the principal roles of SENP3 in the nucleoli is to regulate ribosomal biogenesis, specifically of the 60S ribosomal subunit. Briefly, mammalian 80S ribosomes result from assembly of a large 60S subunit, composed of 5S, 5.8S and 28S rRNAs, and a small 40S subunit comprised of the 18S rRNA. A high number of ribosomal proteins are associated with each subunit. SENP3 is implicated in promoting the maturation of the 28S rRNA by de-SUMOylating several nuclear proteins, including NPM1 (27). The observation that SENP3 was unable to accumulate in the nucleoli during *B. abortus* infection prompted us to investigate if NyxA and NyxB could impact other nucleoli proteins associated with biogenesis of the 60S ribosomal subunit, as SENP3. We did not observe any effect on the nucleolar levels of Pescadillo (PES1) (Supplementary Figure 7A), involved in the maturation of the 28S and 5.8S rRNAs and that interacts with NPM1 for nucleolar targeting but does not interact with SENP3 (29). This suggests that not all nucleolar ribosomal biogenesis associated proteins are impacted during *Brucella* infection. A slight effect on NPM1 nucleolar accumulation was observed in some *B. abortus* infected cells (Supplementary Figure 7B), but to a much lower extent than SENP3, suggesting that the bulk of nucleolar NPM1 remained unaffected.

We next analysed the nucleolar accumulation of the VCP-like AAA-ATPase (NVL), also part of the complex of proteins involved in the maturation of the 28S and 5.8S rRNAs (30). Although significant NVL staining was still detected in the nucleoli of infected cells, we observed a striking cytoplasmic accumulation of NVL in punctate structures in all *B. abortus* infected cells at 48h post-infection that were rarely visible in non-infected cells (Figure 5A). We have named these structures *Brucella*-induced foci (Bif).

**Fig. 5.**
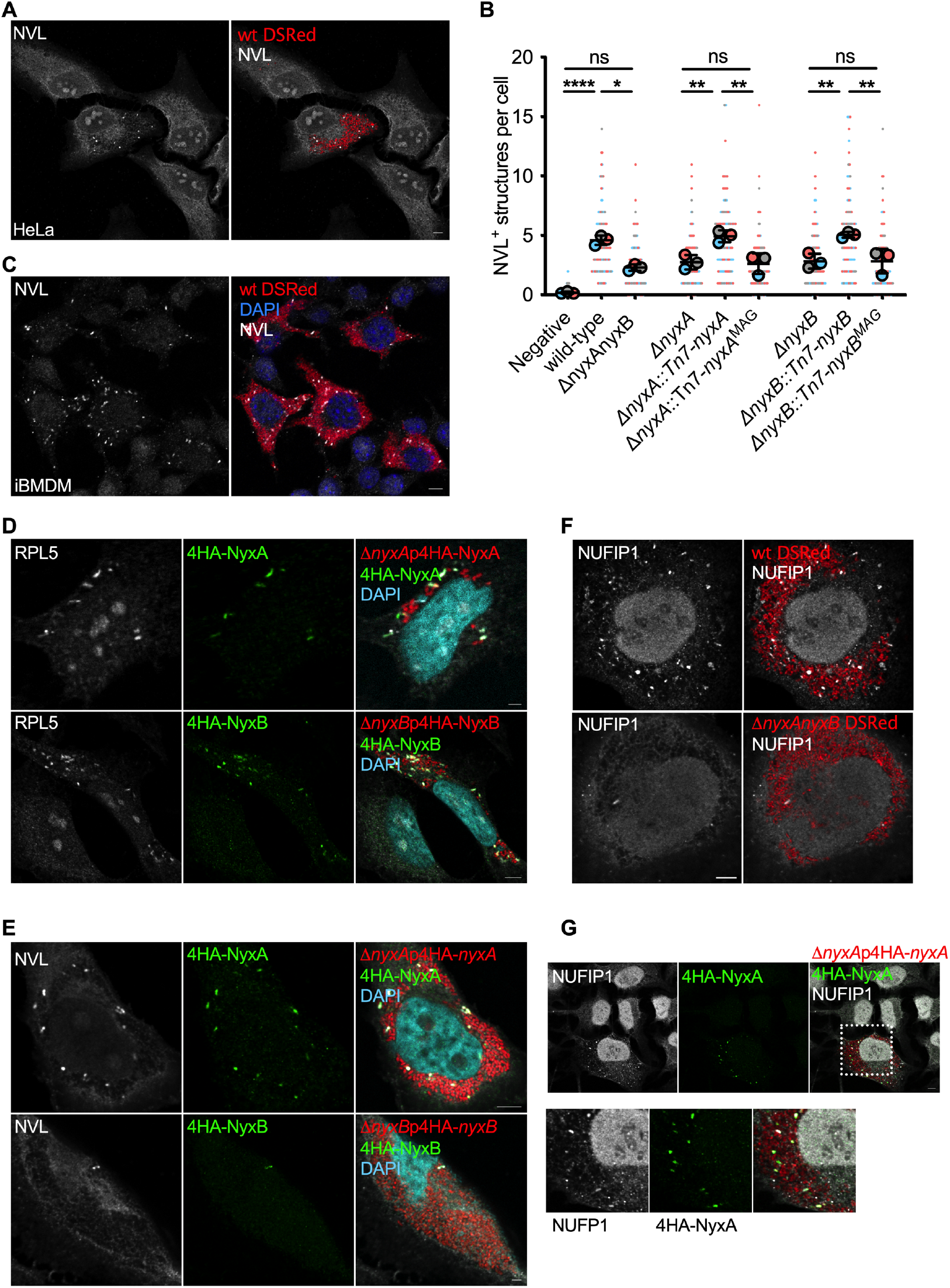
*B. abortus* NyxA and NyxB induce cytoplasmic punctate accumulation of NVL, RPL5 and NUFIP in *Brucella*-induced foci (Bif). **(A)** Confocal microscopy image of wild-type *B. abortus* (expressing DSRed) infected HeLa cells or **(C)** immortalised bone-marrow derived macrophages (iBMDM) labeled with an anti-NVL antibody (white). **(B)** Quantification of the number of NVL-positive cytoplasmic structures in mock infected control cells in comparison to wild-type or a mutant strain lacking each or both *nyxA* and *nyxB*, complemented strains or expressing MAG mutants. Data are represented as means ± 95% confidence intervals from 3 independent experiments. Each experiment is colour coded and all events counted are shown. Data were analysed using one-way ANOVA by including all comparisons with Tukey’s correction. Not all comparisons are shown but are available in Supplementary Table 4. **(D)** Representative confocal microscopy images of HeLa cells infected with either Δ*nyxA* expressing DSRed and 4HA-NyxA (top) or Δ*nyxB* expressing DSRed and 4HA-NyxB for 48h and labelled for RPL5 (white) and DAPI (cyan) or (E) labelled for NVL. **(F)** HeLa cells infected for 48h with wild-type or Δ*nyxAnyxB B. abortus* (DSRed) labeled with an anti-NUFIP1 antibody (white) or **(G)** infected with Δ*nyxA* expressing DSRed and 4HA-NyxA (green). All scale bars correspond to 5 μm.

To determine if the *Brucella* Nyx effectors contributed to the formation of NVL-positive Bif, we quantified their number in HeLa cells infected either wild-type or a mutant lacking both NyxA and NyxB. We found that cells infected with Δ*nyxAnyxB* had fewer NVL-positive Bif (Figure 5B), suggesting the Nyx effectors contribute to their induction. Using single Nyx mutant strains to assess the contribution of each effector we found that both effectors contribute to the formation of NVL-positive Bif (Figure 5B). These differences were not due to different intracellular bacterial numbers as *nyx* mutant strains replicate as efficiently as the wild-type in HeLa cells (Supplementary Figure 4). Importantly, both mutants were fully complemented (Figure 5B). However, expression of either NyxA and NyxB carrying the MAG mutations, that are unable to interact with SENP3, failed to complement the mutant phenotypes (Figure 5B).

To determine if NyxA/B could interact directly with NVL we carried out a pull-down experiment with the purified effector proteins. Unlike what was observed for SENP3, NVL was not found in the eluted samples suggesting it is not part of the same complex (Supplementary Figure 8A). These results indicate that induction of NVL accumulation in cytoplasmic punctate structures during *Brucella* infection is dependent on both NyxA and NyxB and the acidic groove required for SENP3 interaction. To confirm this phenotype, we infected immortalised bone marrow-derived macrophages (iBMDMs), a well-established model of *Brucella* infection. Strong induction of NVL cytoplasmic punctate accumulation was observed in iBMDMs infected with wild-type *B. abortus* (Figure 5C), in a Nyx-dependent manner (Supplementary Figure 8B).

### *Brucella*-induced NVL foci are also enriched in the ribosomal protein RPL5 and the Nyx effectors

Next, we aimed to identify the nature of the cytoplasmic NVL-positive structures formed during *B. abortus* infection. NVL interacts with the ribosomal protein 5 (RPL5), a component of the 60S ribosomal subunit, responsible for transporting NVL to the nucleoli (31). We hypothesised if RPL5 could also be retained in the cytoplasm, and therefore analysed RPL5 distribution in cells infected with wild-type *B. abortus*. We observed the same striking accumulation of RPL5 in cytoplasmic structures in all infected HeLa cells (Supplementary Figure 9A) and bone marrow-derived macrophages (Supplementary Figure 9B). As these structures were reminiscent of the localisation of translocated 4HA-NyxA and NyxB (Figure 1C), we infected cells with strains expressing 4HA-tagged effectors and co-stained for NVL or RPL5. Indeed, RPL5-positive cytoplasmic structures induced upon *Brucella* infection contained NyxA and NyxB (Figure 5D). Similarly, NVL significantly co-localised with the 4HA-tagged Nyx effectors in both HeLa (Figure 5E) and macrophages (Supplementary Figure 10), as well as when using 3Flag-tagged NyxA (Supplementary Figure 11). Due to antibody incompatibility we could not co-label with NVL and RPL5 simultaneously. Nonetheless, these data overwhelmingly support the idea that Bif are enriched in NVL, RPL5, and both Nyx effectors during infection.

### *Brucella* infection induces nucleus to cytoplasm shuttling of ribophagy receptor NUFIP1

Formation of cytoplasmic aggregates has been described in several bacterial infections, including stress granules (32), P-bodies (33) and U-bodies (34). None of these were observed at 48h post-infection (Supplementary Figure 12A and B) when Bif are normally present. Labelling with the FK2 antibody did reveal the presence of aggregates of mono- and poly-ubiquitinated proteins in some *B. abortus* infected cells, but these did not colocalise with NVL (Supplementary Figure 12C). As *Brucella* infection was found to induce the cytoplasmic accumulation of both NVL and RPL5, we focused on other cellular processes known to result in cytoplasmic accumulation of nucleolar proteins. Recent reports have described NUFIP1 as a nucleo-cytoplasmic shuttling protein that accumulates in the cytoplasm upon starvation (8). In this context, NUFIP1 acts as a receptor for ribophagy, a specialised autophagy process dedicated to the degradation of ribosomes to generate nutrients (8). In view of the presence of RPL5 in these structures, a component of the 60S ribosomal subunit we hypothesised that *B. abortus* infection could result in induced ribophagy. Indeed, cells infected with wild-type *B. abortus* showed an increase in cytoplasmic NUFIP1 compared to non-infected cells, with the appearance of bright punctate structures in a manner dependent on NyxA and NyxB (Figure 5F and Supplementary Figure 13A). These structures also contained Nyx effectors (Figure 5G) and were positive for NVL (Supplementary Figure 13B), revealing that Bif also contain NUFIP1.

### NyxA/B-mediated induction of Bif and de-localisation of nucleolar proteins is dependent on Beclin1 and negatively regulated by SENP3

To determine if the presence of NUFIP1 could be related to its function as a ribophagy receptor, we depleted the autophagy-initiation protein Beclin1 which has been shown to be essential for initiation of ribophagy (7). We quantified the number of Bif in wild-type infected cells, using NVL labelling as a marker. We found a reduction in the number of Bif in the absence of Beclin1 (Figure 6A), consistent with a ribophagy-derived process.

**Fig. 6.**
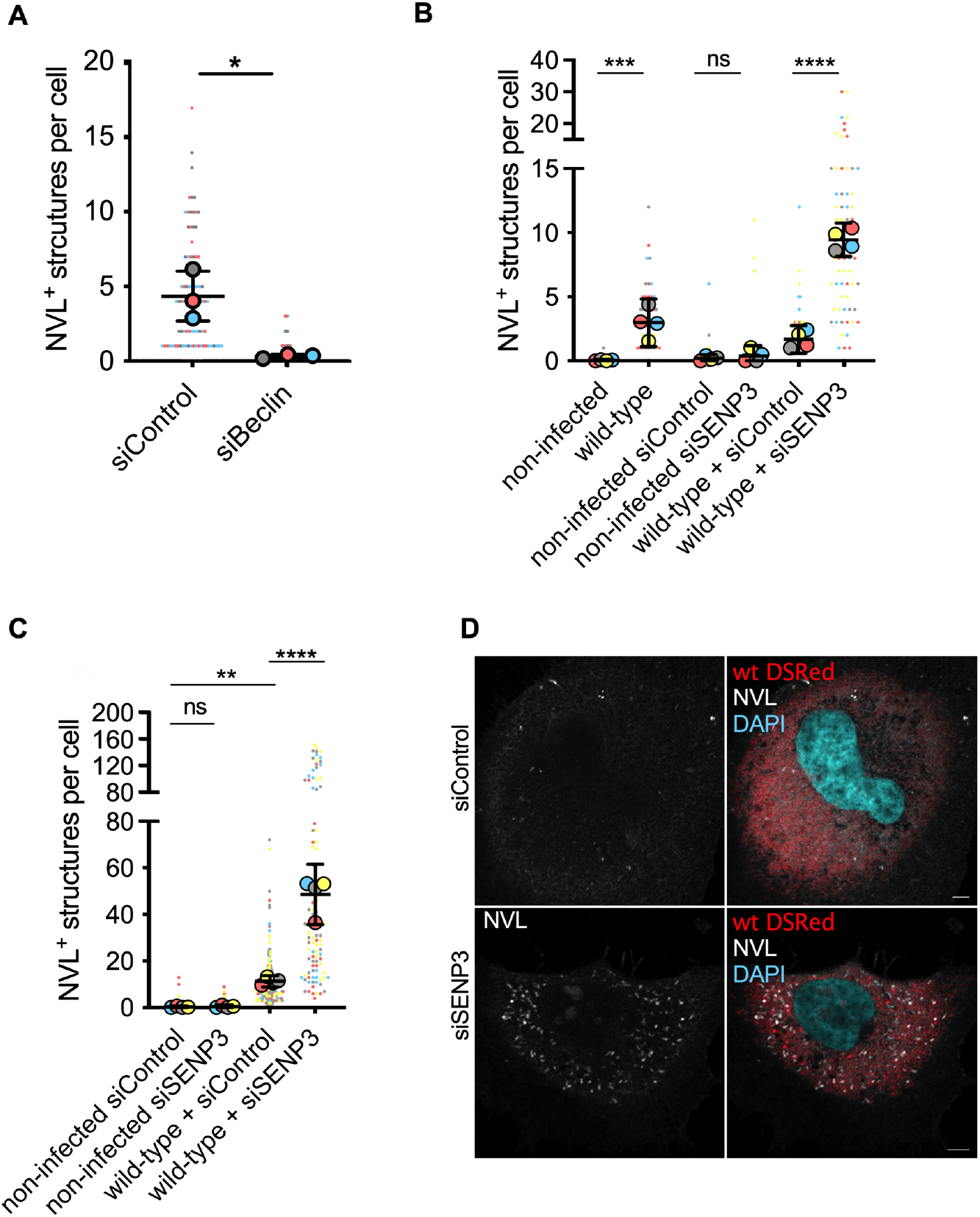
Formation of Bif is dependent on Beclin1 and negatively regulated by SENP3. **(A)** Quantification of the number of NVL-positive structures in cells infected by the wild-type *B. abortus* and depleted for Beclin1 with siRNA or treated with scrambled siRNA (siControl). Data are represented as means ± 95% confidence intervals from 3 independent experiments. **(B)** Quantification of the number of NVL-positive cytoplasmic structures in HeLa cells pre-treated with control siRNA or siSENP3 and infected for an additional 24h with wild-type *B. abortus* or **(C)** 48h. Mock infected cells are included as controls. Data are represented as means ± 95% confidence intervals from 4 independent experiments. Each experiment is colour coded and all events counted are shown. Data were analysed using one-way ANOVA by including all comparisons with Tukey’s correction. Not all comparisons are shown. **(D)** Representative images of NVL cytoplasmic punctate accumulation (white) in HeLa cells treated with control siRNA (top) or depleted for SENP3 (bottom) followed by infection for 48h with *B. abortus* expressing DSRed. All scale bars correspond to 5 μm.

Recently, SENP3 has been shown to de-SUMOylate Beclin1 in the cytoplasm upon starvation and negatively regulate its activity, acting as a switch off mechanism to prevent excessive autophagy (4). NyxA and NyxB interaction with SENP3 and subsequent nuclear mislocalisation is likely to lead to decreased activity of SENP3 elsewhere in the cell. Therefore, to determine if the absence of SENP3 activity could account for the induction of NVL cytoplasmic structures, we depleted SENP3 and infected HeLa cells with wild-type *B. abortus*. The depletion of SENP3 resulted in a substantial rise in cytoplasmic NVL-positive Bif being formed as early as 24h post-infection (Figure 6B). At 48h post-infection, wild-type infected cells showed very high number of Bif in the absence of SENP3, with some cells containing more than 100 puncta (Figure 6C and D). Importantly, depletion of SENP3 alone was not sufficient to induce NVL cytoplasmic accumulation, suggesting this phenotype is triggered by the infection. Therefore, we hypothesised that it could constitute a generalised cellular response to intracellular bacteria. We selected another intracellular pathogen, *Legionella pneumophila*, that multiplies in an ER-derived vacuole as observed for *B. abortus* (35). However, infection with *L. pneumophila* did not result in NVL cytoplasmic accumulation (Supplementary Figure 14), suggesting this phenomenon is specifically induced during *B. abortus* infection.

In summary, these data show that Nyx-mediated accumulation of nucleolar proteins in cytoplasmic structures induced during *Brucella* infection is dependent on Beclin1 and negatively regulated by SENP3.

## Conclusions

Here we report the identification of two new *Brucella* effectors that target SENP3 and induce its delocalisation from the nucleoli during infection. The action of NyxA and NyxB mediate the appearance of cytoplasmic structures enriched in NUFIP1, NVL and RPL5, whose formation is dependent on Beclin1 and negatively regulated by SENP3. We, therefore, propose that NyxA/B interaction with SENP3 leads to enhancement of a ribophagy-like process induced upon *Brucella* infection, that contributes to intracellular proliferation of *Brucella*. Our data point to a key role of SENP3 in the infectious process.

How these effectors are translocated into host cells remains unclear. The TEM-fusion assays indicate that the translocation of NyxA is mostly dependent on the T4SS, unlike NyxB. Our work shows that the two proteins adopt the same ternary and tertiary structures and thus the main differences between these two effectors reside in the N-terminal domain and the last 5 C-terminal residues. A T4SS secretion signal may be present within these regions. Unfortunately, we could not use the 4HA-tagged Nyx effectors to dissect their secretion mechanisms as they were not detected at the early stages of the infection when numbers of wild-type and *virB* mutant bacteria are equivalent. We are currently working on alternative reporter systems with higher sensitivity to better understand the kinetics and mechanisms of translocation, as NyxA and NyxB may constitute an interesting tool to understand the molecular basis of substrate selection by the *Brucella* VirB/D T4SS.

The discovery of NyxA was based on the presence of a potential CAAX motif. However, NyxB, which has the same intracellular location and function, and is more than 80% identical in amino acid sequence, does not contain this putative lipidation site. Therefore, we have no evidence to date that this sequence is important for NyxA function, nor if it constitutes a functional lipidation motif. NyxA does contain the critical conserved cysteine within this motif, followed by the aliphatic amino acid leucine. However, one glutamic acid is also present. Preliminary experiments from our lab show no difference in intracellular localisation of NyxA lacking the C-terminal last four amino acids, consistent with the possibility that it is not necessary for membrane targeting. Furthermore, translocated Nyx effectors were not associated with the vacuole membrane enclosing replicating bacteria nor the plasma membrane. Instead, 4HA-NyxA/B localised to cytoplasmic punctate structures and the nucleus. Interestingly, cytoplasmic foci were detectable at 24h after infection whereas nuclear aggregates were only visible at 48h and onwards, suggesting that cytoplasmic foci precede the nuclear import. However, our transfection data suggest that the presence of the 4HA tag significantly reduces nuclear import. So, it is likely that native untagged NyxA/B are efficiently targeting both nuclear and cytoplasmic compartments during infection.

NyxA/B are two *Brucella* nucleomodulins impacting the sub-nuclear localisation of SENP3 and subsequently of other nucleolar proteins such as RPL5 and NVL. Although both effectors contribute to these phenotypes, deletion of both genes is not sufficient to obtain the phenotype of non-infected cells. These results suggest that additional effectors may be involved in SENP3 delocalisation, either directly or indirectly, which would explain why a mutant lacking both NyxA and NyxB does not have a replication defect. Full depletion of SENP3, on the other hand, results in a very high accumulation of Bif and renders *B. abortus* intracellular replication less efficient. One possible explanation is that accumulation of a too large amount of Bif is harmful to the cell and/or the bacteria. Alternatively, siRNA depletion of SENP3 occurs at an earlier stage of the replication than the effect induced upon NyxA/B translocation.

Although ectopic expression of the Nyx effectors alone is sufficient to delocalise SENP3, it is not enough to induce Bif formation. *Brucella* infection must cause stress or activate a specific signalling pathway that, in the context of the NyxA/B interaction with SENP3, results in the formation of Bif in infected cells. Although *Brucella* is considered a stealthy pathogen, its intracellular trafficking and replication generate several stresses in the cell, including ER stress. Extensive intracellular multiplication of *Brucella* is likely to lead to starvation-like conditions. Indeed, important metabolic switches have been described during infection (36,37). Infected cells have been shown to have a reduced ability to utilise amino acids suggesting intracellular *B. abortus* is competing with the cell for nutrients (36), and additional studies support the notion that *Brucella* is using amino acids from the host (38,39).

A recent study has implicated SENP3 in the regulation of autophagy (4). During starvation conditions, SENP3 is delocalised to the host cytoplasm, where it de-SUMOylates Beclin1, which reduces its interaction with other autophagy components, acting as a negative regulator of autophagy. The levels of Beclin1 SUMOylation could also be controlling ribophagy, where SENP3 would serve as an overflow valve to prevent excessive degradation of the cell’s ribosomes. *Brucella* extensive replication may induce significant stress reminiscent of starvation conditions that would, in turn, induce autophagy. The retention of SENP3 by NyxA and NyxB would prevent this negative regulator from reaching the cytoplasm, de-SUMOylating Beclin1, and switching off autophagy during *Brucella* infection.

The presence of RPL5 and NUFIP1 in Bif and the requirement of Beclin1 for their formation led us to hypothesise that these could be ribophagy-derived. Induction of ribophagy during *Brucella* infection could provide a source of nutrients to the cell and potentially replicating bacteria. Curiously, proteomics studies have shown a reduction in bacterial ribosomal proteins during the early stages of *Brucella* infection (38), which could indicate *Brucella* is using them as a nutrient source. *Brucella* could be eating the cell’s ribosomes at later stages of the infection cycle to sustain massive intracellular replication, but this remains speculative. Additional effectors could have enzymatic activities or recruit specific enzymes contributing to this process. This is nicely illustrated by the effector BPE123 that has been shown to control host metabolism by interacting and recruiting α-enolase to the vacuolar membrane, enhancing its activity, a process essential for intracellular replication (40).

The process of ribophagy is still poorly understood, with additional pathways being implicated in the degradation of ribosomes during starvation, including bulk autophagy, ER-phagy and processes fully independent of autophagy. Indeed, recent studies suggest that during acute nutrient stress, ribosomal degradation predominantly occurs independently of autophagy (41), highlighting the complexity of this process. Although NUFIP1 has been shown to interact with ribosomes in the context of their selective autophagy (8), playing a key role in cell survival, it has also been shown to shuttle from the nucleus to lysosomes upon mechanical stress, without ribophagy induction (42). This is consistent with its participation in multiple stress responses. Therefore, it remains possible that NUFIP1 present in Bif has alternative functions, such as anchoring the 60S ribosomal subunit or in stress response. However, we have found that the formation of Bif requires Beclin1 and, therefore, highly suggestive of an autophagic process that *Brucella* modulates to benefit prolonged intracellular replication. Interestingly, additional nucleolar targets connected to ribosomal biogenesis were identified as potential interacting partners of NyxA in the yeast two-hybrid screen. CEBPZ, also known as Noc1, is involved in both pre-rRNA processing and maturation of pre-ribosomes (43). It has been shown to participate in the intra-nuclear transport of the 60S subunits (43). ARL6IP4 is an RNA splicing regulator shown to interact with core components of the 60S ribosomal subunit (44). It is therefore possible that the Nyx effectors are targeting large complexes of proteins associated with maturation of the 60S ribosomal subunit, retaining them in the vicinity of multiplying bacteria. Alternative, these may simply constitute markers of a ribophagy-derived process elicited by extensive intracellular bacterial multiplication and maintained by the nucleoplasmic retention of SENP3 by NyxA and NyxB.

## Materials and Methods

### Cells, culture conditions and drug treatments

HeLa and RAW cell lines were obtained from ATCC and were grown in DMEM supplemented with 10% of fetal calf serum. Immortalised bone marrow-derived macrophages (45) from C57BL/6J mice were maintained in DMEM supplemented with 10% FCS and 10% spent medium from L929 cells that supplies MC-CSF. All cells were routinely tested and were mycoplasma free. When indicated, cells were treated with 0.5 mM arsenite (Sigma) for 30 min to induce stress granules and P-bodies and treated with 5 μM thapsigargin (Sigma) 4h to induce U-bodies.

### *Brucella* strains, cultures and infections

*B. abortus* 2308 was used in this study. Wild-type and derived strains were routinely cultured in liquid tryptic soy broth and agar. 50 μg/ml kanamycin was added for cultures of DSRed and 50 μg/ml ampicillin for strains expressing pBBR1MCS-4 (46)(4HA and 3Flag constructs) or complemented strains (pUCTminiTn7T_Km) (47). The list of all *Brucella* strains constructed is included in Supplementary Table 5. For infections, eukaryotic cells were plated on glass coverslips 18h before infection and seeded at 2×104 cell/well and 1×105cells/well for 24 and 6 well plates, respectively. Bacterial cultures were incubated for 16h from isolated colonies in TSB shaking overnight at 37 °C. Culture optical density was controlled at 600 nm. Bacterial cultures were diluted to obtain a multiplicity of infection (MOI) of 1:500 for epithelial cells and 1:300 for iBMDM in the appropriate cell culture medium. Inoculated cells were centrifuged at 400 × g for 10 min to initiate bacterial-cell contact followed by incubation for 1h at 37°C and 5% CO_2_ for HeLa cells and 15 min for iBMDMs. Cells were then washed 3 times with DMEM and incubated a further hour with complete media supplemented with gentamycin (50 μg/mL) to kill extracellular bacteria. The gentamycin concentration was then reduced to 10 μg/mL by replacing the media. At the different time points, coverslips were fixed for immunostaining. For enumeration of bacterial colony forming units (CFU), cells were lysed in 0.1% Triton for 5 min and a serial dilution plated in tryptic soy agar.

### *Brucella* expressing vectors

#### TEM vectors

DNA fragments coding for NyxA and NyxB were obtained by PCR amplification from the *B. abortus* 2308 genome, digested with XbaI and cloned into pFlagTEM1 (48). After verification by sequencing, plasmids were introduced into *B. abortus* 2308 or Δ*virB9* by electroporation.

#### Deletion mutants

*B. abortus* 2308 knockout mutant Δ*nyxA* and Δ*nyxB* were generated by allelic replacement. Briefly, upstream and downstream regions of about 750 bp flanking the each gene were amplified by PCR (Q5 NEB) from the *B. abortus* 2308 genomic DNA. An overlapping PCR was used to associate the two PCR products and the Δ*nyxA* or Δ*nyxB* fragments were digested and cloned in a EcoRI digested suicide vector (pNPTS138) (49). These vectors were then electroporated in *B. abortus* and transformants selected using the kanamycin resistance cassette of the pNPTS138 vector. The loss of the plasmid concomitant with either deletion or a return to the wild type phenotype was then selected on sucrose, using the sacB counter selection marker also present on the vector. Deletant (Δ) strain was identified by PCR.

#### Complementing strains

The complementing strain was constructed by amplifying either *nyxA* or *nyxB* and their corresponding promoter region (500 bp upstream) with the PrimeStar DNA polymerase (Takara). Insert and pmini-Tn7 (47) were digested with SacI and BamHI and ligated overnight. Transformants were selected on kanamycin 50 μg/mL and verified by PCR and sequencing. To obtain the complementing strain the Δ*nyxA* or Δ*nyxB* mutants were electroporated with pmini-Tn7-*nyxA* or -*nyxB*, respectively, with the helper plasmid pTNS2. Electroporants were selected with kanamycin 50 μg/mL and verified by PCR.

### 4HA and FLAG vectors

To construct the 4HA tagged vectors, the *nyxA* and *nyxB* genes were sub-cloned in plasmid pMMB 207c (23). The *nyxA* and *nyxB* genes were previously amplified from pGEM-T-easy-*nyxA* or -*nyxB* plasmid. The primers used to amplify *nyxA*/B have KpnI and HindIII restriction sites on the forward and reverse primers, respectively. The PCR products were cleaved by these restriction enzymes to be inserted into the same restriction sites of pMMB207c downstream of the 4HA tag. Subsequently, 4HA-*nyxA* or -*nyxB* were amplified with primers having the SacI and SpeI restriction sites on the forward and reverse primers respectively. The PCR products were cleaved by these restriction enzymes to be inserted into the same restriction sites of pBBR1MCS-4.

### MAG mutants

The mutants Nyx^AMAG^ and NyxB^MAG^ were obtained from pET151-*nyxA* or -*nyxB* and pmini-Tn7-*nyxA* or -*nyxB* using QuickChange Site-Directed Mutagenesis. The mutants NyxA^MAG^ and NyxB^MAG^ correspond to three mutated amino acid residues. These three mutations were introduced one by one sequentially by mutating residues Y62R, D76R, E78R and Y66R, D80R, E82R for NyxA and NyxB, respectively. Resulting vectors were either introduced in *B. abortus*, or *B. abortus* expressing DSRed or GFP. All the primers used are listed in Supplementary Table 6 and all constructs were verified by sequencing.

### Eukaryotic expression vectors

The NyxA and NyxB constructs were obtained by cloning in the gateway pDONRTM (Life Technologies) and then cloned in the pENTRY Myc or HA. The NyxA^MAG^ and NyxB^MAG^ constructs were obtained by site-directed mutagenesis as described above. The 4HA-*nyxA* and 4HA-*nyxB* genes were amplified from the plasmid pBBR1MCS4-4HA-*nyxA* or -*nyxB*. The primers used to amplify 4HA-*nyxA* or -*nyxB* have the BamHI and EcoRI restriction sites in forward and reverse primers, respectively. The PCR products were cleaved by these restriction enzymes to be inserted into the same restriction sites of pcDNA3.1. All the primers used are listed in Supplementary Table 6.

### Bacterial expression vectors

NyxA and NyxB were amplified by PCR and inserted into the pET151D topo vector following the manufacturer’s procedure (Invitrogen) to obtain His-NyxA and His-NyxB. An additional V5 tag is also present. His-NyxAMAG and His-NyxB^MAG^ were obtained by site-directed mutagenesis as described above. For expression and purification of SENP_7-159_, a vector with *E. coli* codon-optimised SENP3 was obtained from Thermofisher and used as a template. SENP_7-159_ was cloned into pRSF-MBP vector. This vector corresponds to pRSFDuet-1 (Novagen) but modified to insert 6xHis-MBP and a protease recognition site for tobacco etch virus (TEV) protease. All the primers used are listed in Supplementary Table 6.

### Immunofluorescence microscopy

At the indicated time points, coverslips were washed twice with PBS, fixed with AntigenFix (MicromMicrotech France) or PFA 3.2% (Electron Microscopy Sciences) for 20 minutes and then washed again 4 times with PBS. To detect translocated HA and Flag-tagged effectors, permeabilisation was carried out with a solution of PBS containing 0.3% triton for 10 minutes to minimise permeabilisation of BCV membranes. This was followed by blocking with PBS containing 2% bovine serum albumin (BSA), 10% horse serum. Primary antibodies diluted in 2% BSA and 10% HS were then incubated at 4 °C overnight in the blocking solution. Subsequently, the coverslips were washed twice in PBS and incubated for 1h with secondary antibodies. Finally, coverslips were washed twice in PBS and once in ultrapure water. Lastly, they were mounted on a slide with ProLongGold (Life Technologies). The coverslips were visualised with a Confocal Zeiss inverted laser-scanning microscope LSM800 and analysed using ImageJ (50) software and final figures assembled with FigureJ (51). Note that neither blocking nor antibody solutions did not contain Triton (only the permeabilisation step). It is important to point out that occasional intravacuolar labeling was observed in weak DSRed labeled bacteria, perhaps indicative of fragilised BCVs. The same protocol was done for NUFIP1. For all other antibodies, when used in the absence of 4HA or 3Flag for translocated effectors, cells were permeabilised with a solution of PBS containing 0.5% triton for 20 minutes followed by blocking for 20 minutes in a solution of PBS containing 2% bovine serum albumin (BSA), 10% horse serum, 0.3% triton. Coverslips were then incubated either at 4 °C overnight with primary antibody diluted in the blocking solution containing 0.3% Triton. Successful labeling was also achieved with 3h incubation at RT. The subsequent steps were identical as described above. The NVL antibody was not stable when stored at −20°C for prolonged periods. All fixations, dilutions and sources for each antibody are indicated in Supplementary Table 7.

### Transfections and siRNA

All cells were transiently transfected using Fugene^®^ (Promega) overnight, according to manufacturer’s instructions. HeLa cells were seeded 18h prior to experiments at 2×104 cells/well for 24 well plates. siRNA experiments were done with Lipofectamine^®^ RNAiMAX Reagent (Invitrogen) according the protocol of the manufacturers. Cells were seeded at 2×104 cells/well. Importantly, siRNA depletion of SENP3 was done by treatment with 5 nM siRNA every day for 72h. Some cell detachment was observed following the 3 days siRNA treatment. Results were confirmed with an alternative siRNA from HorizonDiscovery, efficient after 2 days of silencing. For siBeclin depletion, 2 days of treatment was used with 10 nM. A scrambled siControl was always included in all experiments. All siRNA used are described in Supplementary Table 6.

### *Legionella* infection

*Legionella pneumophila* strain Paris was cultured in N-(2-acetamido)-2-aminoethanesulfonic acid (ACES)-buffered yeast extract broth or on ACES-buffered charcoal-yeast (BCYE) extract agar (52). Human monocyte cell line (THP-1) was cultured and infected as previously described (53). For immunofluorescence analyses cells were fixed in 4% paraformaldehyde, permeabilised with PBS-triton 0.5% and stained with 4-6-diamidino-2-phenylindole (DAPI), Phalloidin and primary anti-NVL antibody (16970-1-AP). Immunosignals were analysed with a Leica SP8 microscope at 63X. Images were processed using ImageJ software.

### Pulldown assays

For pulldown experiments with two purified proteins, 50 μg of His tag recombinant protein was incubated with recombinant protein during 2 h at 4°C, then incubated within gravity flow column (Agilent) containing 80 μl Ni-NTA agarose beads (Macherey-Nagel) during 1 h beforehand washed in water and pre-equilibrated in equilibrium buffer 20 mM Tris– HCl pH7.5, 250 mM NaCl. The column was washed successively three times in equilibrium buffer supplemented with 25 mM imidazole, three times in equilibrium buffer and eluted in equilibrium buffer supplemented with 500 mM imidazole. Proteins eluted were separated by SDS–PAGE, transferred to a PVDF membrane, incubated with specific primary antibodies for 1 h and detected with horseradish peroxidase (HRP)-conjugated secondary antibodies by using ClarityTM Western ECL Blotting Substrate (Bio-Rad). For pulldown assays between cell extract and purified Nyx and MAG mutants, HeLa cells were seeded at 5.10^5^ in 10-cm cell culture dish and incubated overnight in a 37°C humidified atmosphere of 5% CO2. 22h after incubation, cells were washed in ice-cold PBS, harvested and resuspended in 200μl of RIPA buffer (Sigma) supplemented with phenylmethylsulfonyl fluoride (Sigma) and protease inhibitor cocktail (Roche). Extracts were incubated on ice for 15 min with periodical pipetting, then centrifuged at 16,000xg at 4°C for 20 min to pellet cell debris. Supernatant was transferred to a fresh tube. 50 μg of His tag recombinant protein was incubated with 80μl Ni-NTA agarose beads (Macherey-Nagel) within gravity flow column (Agilent) during 1 h beforehand washed in water and pre-equilibrated in equilibrium buffer 20 mM Tris– HCl pH7.5, 250 mM NaCl. The cell extract was incubated within the column (containing His tag recombinant protein and Ni-NTA agarose beads) during 1h at 4°C under agitation. The column was washed successively three times in equilibrium buffer supplemented with 25 mM imidazole, three times in equilibrium buffer and eluted in equilibrium buffer supplemented with 500 mM imidazole (54).

### Quantification of SENP3 localisation

Colocalisation analysis for the transfected HeLa cells were performed with a custom ImageJ (50)/Fiji (55)-based macro (http://c2bp.ibcp.fr/CL_ArthurL_coloc_2-corr.ijm), that segmented the nuclei and the nucleoli of the cells in each image, classified the cells in two classes according to the intensity of HA-NyxA/NyxB, and then measured in the areas of each nucleus the Pearson correlation coefficients (by calling the plugin Coloc2 - https://github.com/fiji/Colocalisation_Analysis) between the signal of SENP3 and the nucleolin as well as between the signals of SENP3 and HA-NyxA/NyxB. For each nucleus, the ratio between the mean intensity of the SENP3 signal in the nucleoli and the mean intensity of the SENP3 signal outside the nucleoli is also calculated. The details can be found in the source code and the comments of the macro. For the *Brucella* infected HeLa cells, a pipeline in the software CellProfiler (56) was used to measure the Pearson correlation coefficient between the signal of SENP3 and the nucleolin, in the nuclei of the cells. The cells were also classified in two classes according to the intensity of the *Brucella* DSRed signal in the perinuclear area. The pipeline can be downloaded here http://c2bp.ibcp.fr/arthur2.cpproj.

### TEM1 translocation assay

RAW cells were seeded in a 96 well plates at 1×10^4^ cells/well overnight. Cells were then infected with an MOI of 500 by centrifugation at 4 °C, 400 g for 5 min and 1 at 37 °C 5% CO2. Cells were washed with HBSS containing 2.5 mM probenicid. Then CCF2 mix (as described in the Life Technologies protocol) and probenicid were added to each well, and incubated for 1.5 h at room temperature in the dark. Cells were finally washed with PBS, fixed using 3.2% PFA and analysed immediately by confocal microscopy (Zeiss LSM800).

### Thermophoresis

The experiments were performed at 20°C on a Monolith NT.115 (Nanotemper) with Premium Coated (Nanotemper) capillaries. The tested interactions were performed in triplica. The labelling of NyxA was performed by covalent coupling of the lysines of the protein to a fluorophore using the RED-NHS (Nanotemper) Protein Labeling Kit. Unlabeled NyxB was used at the following increasing concentrations: 6.67 nM, 13.34 nM, 26.68 nM, 53.3 nM, 106.72 nM, 213.44 nM, 426.88 nM, 853.76 nM, 1.7 μM, 3.4 μM, 6.8 μM, 13.6 μM, 27.2 μM, 54.4 μM, 108.8 μM, 217.6 μM.

### Protein expression and purification

*E. coli* BL21-DE3-pLysS bacteria were transformed with the expression vectors, grown in lysogenic broth (LB) media (Sigma-Aldrich) to OD_280_=0.6 and expression was induced with 1mM IPTG at 37°C for 3 hours. Cells were After harvested by 15 min of centrifugation at 5,000 g and resuspended in lysis buffer 20mM Tris pH 8, 150mM NaCl, 5% Glycerol, 0,1% Triton. Disruption cell was achieved with sonication after addition of antiprotease EDTA-Free cocktail (Roche) and 30U/ml benzonase (Sigma-Aldrich). Cell debris were removed by centrifugation 30 min at 20,000g at 4°C. Recombinant protein was purified by chromatography using a Nickel-loaded Hitrap Chelating HP column (GE Healthcare). Unbound material was extensively washed using Tris 20mM pH8, NaCl 300mM, 25 mM Imidazole, 5mM β-mercaptoethanol, 10% Glycerol. An additional washing step with 2 column volumes of 1M NaCl was done before elution of NyxB over a 25 to 500 mM gradient of imidazole over 8 column volumes. Peak fractions were pooled and the His tag was cleaved with TEV protease (500 μg/20mg of eluted protein) in presence of 1 mM DTT and 0,5 mM EDTA in overnight dialysis buffer 20mM Tris pH 8, 150mM NaCl. NyxB was further purified by size exclusion chromatography (Superdex 200 HiLoad 16/600, GE Healthcare) equilibrated in 20mM Tris pH 8, 150mM NaCl, 5% Glycerol. Purity of the sample was assessed by SDS-PAGE. Freshly purified NyxB was concentrated to 21 mg/ml on 3 KDa Amicon Ultra concentrators (Millipore). SeMet-NyxB was produced in M9 minimum medium and purified as above, to a final concentration of pure SeMet-NyxB of 24 mg/ml.

### Crystallisation and data collection

Screening was conducted using a Mosquito workstation (TTP Labtech) on commercial crystallisation solutions with the sitting-drop vapour diffusion technique, against a protein solution. All crystallisation trials were performed at 19°C and visualised on RockImager 182 (Formulatrix). Crystals of native NyxB were obtained with 21 mg/ml NyxB in 25% PEG4000, 0,2M CaCl_2_, 100 mM Tris pH 8. Crystals were frozen in reservoir solution supplemented with 15% Gly. Diffraction data was collected at the European Synchrotron Radiation Facility (ESRF) Beamline line ID23EH1 and crystals diffracted up to 2.5 Å resolution in space group P6_1_22, with 12 molecules per asymmetric unit. Crystals of Se-Met NyxB were obtained using a reservoir solution containing M NaFormate and crystals were cryoprotected in 2.8 M Naformate supplemented with 10% glycerol. Data were collected at SOLEIL on beamline Proxima-2 from a single crystal that diffracted up to 3.7 Å resolution and belonged to the space group P6_2_22, with 2 molecules per asymmetric unit. Diffraction data were processed using XDS (57) and with SCALA from the CCP4 program suite (58). Data collection statistics are indicated in Supplementary Table 2.

The structure was solved using the single anomalous dispersion method on Se-Met crystals using AutoSol from the Phenix program suite (59). An excellent experimental electron density map enabled us to manually build an initial model. The resulting model was then used for molecular replacement with data from native NyxB crystal using Phaser (59). Twelve monomers were positioned and the resulting electron density map was then subjected to the AutoBuild program, part of the Phenix program suite. Model building were completed with sessions of manual model building using Coot combined with model refinement using Phenix (60). The final model was refined to a final Rwork/Rfree of 0.20/0.24 with excellent geometry (Supplementary Table 2). The coordinates and structure factors of NyxB have been deposited in the protein DataBank with the code 7AD4. Figures were generated with Pymol, Molecular Graphics System, Version 2.0 Schrödinger, LLC.

### Size exclusion Multi-angle light scattering

Size exclusion chromatography (SEC) experiments coupled to multi-angle laser light scattering (MALS) and refractomestry (RI) were performed on a Superdex S200 10/300 GL increase (GE Healthcare) for NyxA and a Superdex S75 10/300 GL (GE Healthcare) for NyxB. Experiments were performed in buffer 20 mM Tris pH 8, 150 mM NaCl, 5% Glycerol. 100 μl of proteins were injected at a concentration of 10 mg/ml. On-line MALS detection was performed with a miniDAWN-TREOS detector (Wyatt Technology Corp., Santa Barbara, CA) using a laser emitting at 690 nm and by refractive index measurement using an Optilab T-rex system (Wyatt Technology Corp., Santa Barbara, CA). Weight averaged molar masses (Mw) were calculated using the ASTRA software (Wyatt Technology Corp., Santa Barbara, CA).

### Small-angle X-ray scattering

SAXS data were collected for NyxA and NyxB on BioSAXS beamline BM29, ESRF using an online size-exclusion chromatography setup. 50 μl of protein (10 mg/ml) were injected into a size-exclusion column (Agilent BioSec-3) equilibrated in 50 mM Tris, pH 8.0, 200 mM NaCl. Images were acquired every second for the duration of the size-exclusion run. Buffer subtraction was performed by averaging 20 frames either side of the peak. Data reduction and analysis was performed using the BsxCuBE data collection software and the ATSAS package (61). The program AutoGNOM was used to generate the pair distribution function (P(r)) and to determine Dmax and Rg from the scattering curves (I(q) *versus* q) in an automatic, unbiased manner. Theoretical curves from the models were generated by FoXS (62). *Ab initio* modelling was performed with GASBOR (63).

### Yeast two-hybrid

NyxA was cloned into pDBa vector, using the Gateway technology, transformed into MaV203 and used as a bait to screen a human embryonic brain cDNA library (Invitrogen). Media, transactivation test, screening assay and gap repair test were performed as described (64–66).

### Graphs and stats

For microscopy counts of individual cells, Super Plots were used (67). All data sets were tested for normality using Shapiro-Wilkinson test. When a normal distribution was confirmed we used a One-Way ANOVA test with a Tukey’s correction for multiple comparisons. For two independent variables, a Two-Way ANOVA test was used. For data sets that did not show normality, a Kruskall-Wallis test was applied, with Dunn’s correction. All analyses were done using Prism Graph Pad 7.

## ACKNOWLEDGEMENTS

We would like to thank Rémi Lagorce for help with the purification of the N-terminus of SENP3, Inès Bordin for the optimisation of the expression conditions of NyxB, Marie-Pierre Candusso and Mégane Guinot from Protein Science Facility of SFR Biosciences for help with protein purification of NyxB and crystallography experiments. We are grateful to Gunnar Schroeder (Queen’s University of Belfast) for the pMMB 207c, Renée Tsolis (University of California, Davis) for the pFlagTem, Xavier de Bolle (University of Namur) for the pNPTS138 and Jean Celli (Washington State University) for the pUCTminiTn7T_Km. The LAMP1 and Myc antibodies were developed by Bishop, J.M. were obtained from the Developmental Studies Hybridoma Bank, created by the NICHD of the NIH and maintained at the University of Iowa. We thank the Institut Pasteur and ANR-10-LABX-62-IBEID for the *Legionella* experiments. We acknowledge the contribution of the SFR Biosciences (UAR3444/CNRS, US8/Inserm, ENS de Lyon, UCBL) facilities: Protein Science Facility for mass spectrometry and crystallography experiments and the Plateau Technique Imagerie/Microcopie (PLATIM) for all microscopy studies. We thank staff scientists from beamlines Proxima-2 (synchrotron SOLEIL), ID23EH1 (ESRF) and BM29 (ESRF) for assistance with data collection. These effectors were discovered under the ERA-Net Pathogenomics grant and the remaining work funded by ANR-15-CE15-0011-01. We also thank Steve Garvis and Stéphane Méresse for critical reading of the manuscript. Jean-Paul Borg is a scholar of the Institut Universitaire de France.

## Author contributions

A Louche carried the vast majority of the experimental work with *Brucella* (cell biology, infection studies and biochemistry). A Blanco did the assessment of *Brucella* replication with siSENP3 and discovered NVL/RPL5 delocalisation and Supp Fig 13. She carried out the optimisation of the protein purification of NyxA for the MALS. TLS Lacerda discovered these effectors and showed their translocation and nuclear targeting in transfection. C Lionnet coded the plugIns for quantification of SENP3 and assisted with image analysis. C Bergé performed and analysed SAXS experiments, M Rolando and C Buchrieser the *Legionella* experiment, F Lembo and JP Borg did the Y2H. This work was started in the lab of JP Gorvel who contributed to the discovery of this effector. M Nagahama provided tools and input regarding NVL, RPL5 and ribophagy. V Gueguen-Chaignon and L Terradot supervised and performed the MALS, the crystallography screens and the diffraction data collection. L Terradot solved, analysed the structure of NyxB, designed the MAG mutant and wrote the structural part of the manuscript. SP Salcedo directed, funded and supervised the vast majority of the work, and wrote the manuscript with A Louche. All other authors read and approved the manuscript.

## Notes

### Competing Interest Statement

The authors have declared no competing interest.

