## Supplementary information for "The unique *Brucella* effectors NyxA and NyxB target SENP3 to modulate the subcellular localisation of nucleolar proteins"

### 1. SUPPLEMENTARY RESULTS

Data indicates that the Rg of NyxB and NyxA are 2.9 and 2.7 nm, with $D_{max}$ values of 10.2 nm and 9.6 nm, respectively. These data are in agreement with a dimeric form of the proteins given that theoretical Rgs for dimer 1 and 2 are 2.4 nm and 2 nm, respectively while Rg of a monomer of NyxB is 1.4 nm. The higher Rg observed in solution might be due to the disordered region present at the N-terminal portion of each NyxB monomer (residues 1-16) and NyxA (residues 1-12). Comparison of NyxB experimental SAXS data with theoretical curves obtained with NyxB dimer 1 or dimer 2 indicates that the best fit is obtained with dimer 1 ($\chi^2$= 1.3) compared to dimer 2 ($\chi^2$= 3.3) (Supplementary Figure 4C and D). Ab initio modelling using SAXS data confirmed that the envelope obtained fits better the NyxB dimer1. Similar results were obtained using NyxA (Supplementary Figure 4C and D). Collectively, X-ray, MALS and SEC SAXS data clearly establish that dimer 1 (Figure 3A) is the conformation of NyxB and NyxA in solution. This NyxB dimer relies on reciprocal hydrophobic and electrostatic interactions between $\alpha$4 of one sububnit and $\alpha$6 of the other subunit and between the two $\alpha$4-$\beta$4 loops, burying a total of 530 Å$^2$ (Figure 3A).

### 2. SUPPLEMENTARY TABLES AND FIGURES

**Supplementary Table 1. Table of identified baits for NyxA.**

| Baits from Y2H for NyxA interaction partner | Number of hits |
|---|---|
| Homo sapiens SUMO1/sentrin/SMT3 specific peptidase 3 (SENP3) | 5 |
| Homo sapiens complement component 1, r subcomponent (C1R) | 8 |
| Homo sapiens canopy 4 homolog (zebrafish) (CNPY4) | 2 |
| TAF6 TAF6 RNA polymerase II, TATA box binding protein (TBP)-associated factor | 1 |
| CEBPZ CCAAT/enhancer binding protein (C/EBP), zeta [ Homo sapiens ] | 3 |
| ARL6IP4 ADP-ribosylation-like factor 6 interacting protein 4 | 3 |
| Glutathione S-transferase kappa 1 | 1 |
| Homo sapiens chromosome 11 genomic scaffold, alternate assembly HuRef SCAF_1103279188392<br>Homo sapiens chromosome 11 genomic scaffold, alternate assembly CHM1_1.0<br>Homo sapiens chromosome 11 genomic contig, GRCh37.p10 Primary Assembly | 1 |
| Heparan sulfate proteoglycan 2 (HSPG2) | 1 |
| Homo sapiens RNA binding motif, single stranded interacting protein 3 (RBMS3) | 1 |
| Homo sapiens alkaline phosphatase, intestinal (ALPI), mRNA<br>Homo sapiens alkaline phosphatase, placental-like 2 (ALPPL2), mRNA<br>Homo sapiens alkaline phosphatase, placental (ALPP), mRNA | 1 |

**Supplementary Table 2. Data collection and refinement statistics**

Statistics for the highest-resolution shell are shown in parentheses.

| | Se-Met | Native |
|---|---|---|
| Wavelength (Å) | 0.97930 | 0. 97242 |
| Resolution range (Å) | 48.94 - 3.7 (3.9 - 3.7) | 48.69- 2.5 (2.64 - 2.5) |
| Space group | P 62 2 2 | P 61 2 2 |
| Cell parameters a, b, c (Å) | 77.60 77.60 195.08 | 133.91 133.91 389.51 |
| Total reflections | 101555 (15006) | 1420875 (194308) |
| Unique reflections | 4203 (590) | 72440 (10381) |
| Multiplicity | 24.2 (25.4) | 19.6 (18.7) |
| Completeness (%) | 100.0 (98.6) | 100.00 (100.00) |
| Mean I/sigma(I) | 17.4 (4.4) | 12.5 (2.9) |
| Wilson B-factor | 120.0 | 35.9 |
| R-merge | 0.133 (0.805) | 0.164 (1.035) |
| R-meas | 0.138 | 0.173 |
| CC1/2 | 0.999 (0.972) | 0.996 (0.958) |
| **Refinement** | | |
| R-work | | 0.20 (0.26) |
| R-free | | 0.24 (0.32) |
| Macromolecules | | 11352 |
| Ligands | | 24 |
| Water | | 326 |
| Protein residues | | 1421 |
| RMS (bonds) | | 0.008 |
| RMS (angles) | | 1.12 |
| Ramachandran favored (%) | | 98.50 |
| Ramachandran allowed (%) | | 1.43 |
| Ramachandran outliers (%) | | 0.07 |
| Clashscore | | 7.87 |
| Average B-factor | | 53.9 |
| macromolecules | | 54.0 |
| ligands | | 72 |
| solvent | | 49.3 |

**Supplementary Table 4. Statistical analysis of Figure 5B.**

| Tukey's multiple comparisons test | Summary | Adjusted P Value |
|---|---|---|
| Negative vs. wild-type | **** | <0,0001 |
| Negative vs. ΔnyxAnyxB | ** | 0,0081 |
| wild-type vs. ΔnyxAnyxB | ** | 0,0036 |
| wild-type vs. Δ*nyxA* | * | 0,0247 |
| wild-type vs. Δ*nyxA:Tn7-nyxA* | ns | 0,9984 |
| wild-type vs. Δ*nyxA*:Tn7-*nyxA*$^{MAG}$ | * | 0,0147 |
| wild-type vs. Δ*nyxB* | * | 0,0353 |
| wild-type vs. Δ*nyxB:*Tn7-*nyxB* | ns | 0,9862 |
| wild-type vs. Δ*nyxB:*Tn7-*nyxB*$^{MAG}$ | * | 0,0353 |
| Δ*nyxA* vs. Δ*nyxA:*Tn7-*nyxA* | ** | 0,006 |
| Δ*nyxA* vs. Δ*nyxA*:Tn7-*nyxA*$^{MAG}$ | ns | >0,9999 |
| Δ*nyxA*:Tn7-*nyxA* vs. Δ*nyxA*:Tn7-*nyxA*$^{MAG}$ | ** | 0,0035 |
| Δ*nyxB* vs. Δ*nyxB:*Tn7-*nyxB* | ** | 0,0051 |
| Δ*nyxB* vs. Δ*nyxB:*Tn7-*nyxB*$^{MAG}$ | ns | >0,9999 |
| Δ*nyxB*:Tn7-*nyxB* vs. Δ*nyxB:*Tn7-*nyxB*$^{MAG}$ | ** | 0,0051 |

**Supplementary Table 5. *Brucella* strains used in this study.**

| **Strain name** | **Description** | **Genetic features** | **Resistance** |
|---|---|---|---|
| *Brucella abortus* 2308 | Wild-type (obtained from X. de Bolle) | - | Nalidixic acid natural resistance |
| *B. abortus* DSRed | | pTn7-DSRed | Kanamycin |
| Δ*virB9* | Deletion of *virB9* | | |
| Wt p*bla:nyxA* | TEM1 translocation | pFlagTEM1-NyxA | Chloramphenicol |
| Δ*virB9* p*bla:nyxA* | TEM1 translocation | pFlagTEM1-NyxA | Chloramphenicol |
| Wt p*bla:nyxB* | TEM1 translocation | pFlagTEM1-NyxB | Chloramphenicol |
| Δ*virB9* p*bla:nyxB* | TEM1 translocation | pFlagTEM1-NyxB | Chloramphenicol |
| Wt p*bla:BAB1_0466* | TEM1 translocation | pFlagTEM1-BAB1_0466 | Chloramphenicol |
| Δ*nyxA* | Deletion of *BAB1_0296 (nyxA)* | | |
| Δ*nyxB* | Deletion of *BAB1_1101 (nyxB)* | | |
| Δ*nyxAnyxB* | Deletion of both | | |
| Δ*nyxA:Tn7-nyxA* | Complemented strain | pTn7-*nyxA* | Kanamycin |
| Δ*nyxA:Tn7-nyxA*$^{MAG}$ | Complemented strain with mutated acidic groove (MAG) | pTn7-*nyxA*$^{MAG}$ | Kanamycin |
| Δ*nyxA:Tn7-nyxB* | Complemented strain | pTn7-*nyxB* | Kanamycin |
| Δ*nyxB:Tn7-nyxB*$^{MAG}$ | Complemented strain with mutated acidic groove (MAG) | pTn7-*nyxB*$^{MAG}$ | Kanamycin |
| Δ*nyxA*p4HA-NyxA | Effector imaging | pBBR1MCS-4-4HA-*nyxA* | Ampicilin |
| Δ*nyxA*p3Flag-NyxA | Effector imaging | pBBR1MCS-4-3Flag-*nyxA* | Ampicilin |
| Δ*nyxB*p4HA-NyxB | Effector imaging | pBBR1MCS4-4HA-*nyxB* | Ampicilin |

**Supplementary Table 6. Primers and siRNA used in this study.**

| | | Sequence 5'-3' |
|---|---|---|
| pNPTS138-*nyxA* | Fw1 | GATAATGACGCTAGCTTTCA |
| | Rev1 | GGCCACCCGACAAGCAGTTTGGAGGTTTACCTTTTGTTGA |
| | Fw2 | TCAACAAAAGGTAAACCTCCAAACTGCTTGTCGGGTGGCC |
| | Rev2 | GGGTGTCCTTGAAACATCCA |
| pNPTS138-*nyxB* | Fw1 | TTGGATCAATCCGGCGTGTG |
| | Rev1 | GGGCTTCAACTTCTTTAACCGGCTATTCTTCCTGTCAATT |
| | Fw2 | AATTGACAGGAAGAATAGCCGGTTAAAGAAGTTGAAGCCC |
| | Rev2 | CGTAAACGCTTCGGCAGGGA |
| pFlagTEM1-*nyxA* | fw | TAAGCATTGGTCTAGAATGAACGCTCACACAAACATAA |
| | rev | ACTGCAGTTATCTAGATCAAAGCTCCAAGCATCTAATT |
| pFlagTEM1-*nyxB* | fw | GAGATAGGTGCCTCACTGATTAAGCATTGGTCTAGAATGAACACGCAAGCAACAATA |
| | rev | GTGTGCTGGAATTCGCCCTTACTGCAGTTATCTAGATCAAGGCATCTCGATAAG |
| pFlagTEM1-BAB1_0466 | fw | TAAGCATTGGTCTAGAATGAAAATGTGGACCCTTGC |
| | rev | ACTGCAGTTATCTAGATCACTGTTCTACGCAGCTTA |
| pTn7-*nyxA* | fw | AAAAAAGAGCTCTAAGTGTCTGCCATAGCCGACG |
| | rev | AAAAAGGATCCTCAAAGCTCCAAGCATCTAATTTC |
| pTn7-*nyxB* | fw | AAAAAAGAGCTCTTGGATCAATCCGGCGTGTGC |
| | rev | AAAAAGGATCCTCAAGGCATCTCGATAAGGC |
| pMMB 207c-*nyxA* | fw | AAAAAAGGTACCAACGCTCACACAAACATAAGTGG |
| pMMB 207c-*nyxA* | rev | AAAAAAAAAGCTTTCAAAGCTCCAAGCATCTAATTTC |
| pMMB 207c-*nyxB* | fw | AAAAAAGGTACCAACACGCAAGCAACAATAGATACAGC |
| pMMB 207c-*nyxB* | rev | AAAAAAAAAGCTTTCAAGGCATCTCGATAAGGC |
| pBBRMCS4 4HA-*nyxA* | fw | AAAAAAGAGCTCAAGGAGATATACATATGTACC |
| | rev | AAAAAAACTAGTTCAAAGCTCCAAGCATCTAATTTC |
| pBBRMCS4 3Flag-*nyxA* | fw | TATTCCCGGGGGATCCATGGGTAAGCCTATCCCTAACCCTCTCCTCGGTCTCGATT<br>CTACGAACGCTCACACAAACATAAG |
| | rev | TATAGGGCGAATTGGAGCTCAAGCTCCAAGCATCTAATTT |
| pBBRMCS4 4HA-*nyxB* | fw | AAAAAAGAGCTCAAGGAGATATACATATGTACC |
| | rev | AAAAAAACTAGTTCAAGGCATCTCGATAAGGC |
| pDONOR-*nyxA* | fw | GGGGACAAGTTTGTACAAAAAAGCAGGCTTCAACGCTCACACAAACATAAG |
| | rev | GGGGACCACTTTGTACAAGAAAGCTGGGTCCTAAAGCTCCAAGCATCTAATTTC |
| pDONOR-*nyxB* | fw | GGGGACAAGTTTGTACAAAAAAGCAGGCTTCAACACGCAAGCAACAATAGATA |
| | rev | GGGGACCACTTTGTACAAGAAAGCTGGGTCCTAAGGCATCTCGATAAGGCGGATT |
| pcDNA3.1-4HA-*nyxA* | Fw | GCGTTTAAACTTAAGCTTGGTACCGAGCTCGGATCCAAGGAGATATACATATGTACC |
| | Rev | CGAGCGGCCGCCACTGTGCTGGATATCTGCAGAATTCTCAAAGCTCCAAGCATCTAATTTC |
| pcDNA3.1-4HA-*nyxB* | Fw | GCGTTTAAACTTAAGCTTGGTACCGAGCTCGGATCCAAGGAGATATACATATGTACC |
| | rev | CGAGCGGCCGCCACTGTGCTGGATATCTGCAGAATTCTCAAGGCATCTCGATAAGGC |
| pET151His-*nyxA* | fw | CACCATGAACGCTCACACAAAC |
| | rev | TCAAAGCTCCAAGCATCT |

| | | |
|---|---|---|
| pET151His-*nyxB* | fw | CACCATGAACACGCAAGCAAC |
| | rev | CATTATGCTCCCCTGTTGT |
| NyxA Y62R | fw | CGATTGGCGACCTGCCGCCTATGATG |
| | rev | CGGCAGGTCGCCAATCGTAGCAGTCGAAG |
| NyxA D76R | fw | CCATGAAACGACGGGAACTGATCCAATACG |
| | rev | TTCCCGTCGTTTCATGGCGTTGCCTTC |
| NyxA E78R | fw | CGACGGAGACTGATCCAATACGAAGAGTGGTG |
| | rev | GGATCAGTCTCCGTCGTTTCATGGCG |
| SENP3 $_{7\text{-}159}$ | Fw | CACCATGGCCGGCACCGGTAGCTGGGGTCCGGAACC |
| | rev | TTTGGATCCTTATTTGCTATACAGCAGCATACGAAATGC |
| NyxB D80R | fw | CCATGAAACGACGGGAACTGATCCAATATGAAG |
| | rev | GTTCCCGTCGTTTCATGGCGTTGCCTTC |
| NyxB E82R | fw | AACGACGGAGACTGATCCAATATGAAGATTGGTGC |
| | rev | GGATCAGTCTCCGTCGTTTCATGGCGT |
| NyxB Y66R | fw | CGACTGGCGACCCGCCGCCTATGACGAC |
| | rev | GCGGGTCGCCAGTCGTAGGTGTCGAAGAAGAAGC |

**Supplementary Table 7. Antibodies used in this study.**

| Antibody | Species | Source | Reference | Dilution | Fixation |
|---|---|---|---|---|---|
| SENP3 | Rabbit | Cell Signalling | 5591 | 1/400 | PFA |
| NVL | Rabbit | Proteintech | 16970-1-AP Batch 00008428 | 1/25 | Antigenfix |
| NVL | Mouse | Sigma | HPA028224 | 1/50 | Antigenfix |
| RPL5 | Rabbit | From M. Nagahama lab | | 1/100 | Antigenfix |
| PES1 | Rabbit | ATLAS antibodies | HPA040210 | 1/100 | Antigenfix |
| NPM1 | Rabbit | Abcam | Ab37659 | 1/400 | PFA |
| Nucleolin | Mouse | Invitrogen | 39-6400 | 1/100 | Antigenfix |
| HA | Rat | Roche | #11867423001 clone 3F10 | 1/100 | PFA/Antigenfix |
| Flag | Mouse | Sigma | F1804 (clone M2) | 1/2000 | Antigenfix/methanol |
| Myc | Mouse | DSHB | Clone E10 | 1/1000 | PFA/Antigenfix |
| NUFIP1 | Rabbit | Proteintech | 12515-1-AP | 1/50 | Antigenfix |
| LAMP1 | Mouse | DSHB | Clone H4A3 | 1/200 | PFA/Antigenfix |
| FK2 | Mouse | Enzo | BML-PW8810 | 1/1000 | Antigenfix |
| g3bp (H10) | Mouse | Santa Cruz | sc-365338 | 1/25 | PFA |
| eiF3n (c-5) | Mouse | Santa Cruz | sc-137214 | 1/50 | PFA |
| Xrnl (C-1) | Mouse | Santa Cruz | sc-165985 | 1/25 | PFA |
| DDX20 (Gemin3-12H12) | Mouse | Santa Cruz | sc-57007 | 1/50 | PFA |
| LC3 | Mouse | Nanotools | 0231-100/LC3-5F10 | 1/1000 | Methanol |
| Western blot | | | | | |
| Actin | Mouse | Sigma | A4700 | 1/1000 | |
| His | Mouse | Sigma | H1029 | 1/3000 | |
| V5 | Mouse | Invitrogen | R960-25 | 1/1000 | |
| SENP3 | Rabbit | Cell Signalling | 5591 | 1/400 | |
| NPM1 | Rabbit | Abcam | Ab37659 | 1/400 | |
| Histone 3 | Rabbit | Abcam | Ab8895 | 1/500 | |
| Secondary antibodies | | | | | |
| Donkey anti-mouse AlexaFluor488/555/647 | | Invitrogen | A-21202/A-31570/A-31571 | 1/1000 | |
| Donkey anti-rabbit AlexaFluor488/555/647 | | Invitrogen | A-21206/A-31572/A-31573 | 1/1000 | |
| Donkey anti-rat AlexaFluor488/555/647 | | Invitrogen | A-21208/A-21434/A-21472 | 1/1000 | |

**A**

```
NyxA   1 MNAHTNISGSAA----FDPNAWHHSQMTIREAIDLSQSGGHPYSSPNVPK  46
         ||....|..:|.    ||||||||||||..|||:||:|||||||||||||
NyxB   1 MNTQATIDTAAVAPLNFDPNAWHHSQMTTLEAIELSRSGGHPYSSPNVPK  50

NyxA  47 GFNTVVGFFFDCYDWYPAAYDDEEGNAMKDRELIQYEEWCAKYARKLGLE  96
         |||||||||||.|||||||||||||||||||||||||||:|||||||.||||
NyxB  51 GFNTVVGFFFDTYDWYPAAYDDEEGNAMKDRELIQYEDWCAKYARTLGLE 100

NyxA  97 VKEVEAPAALKVHGIMTLKAYPEALLEIRCLEL-  129
         ||||||||||||||||.||||||||||||.:|:
NyxB 101 VKEVEAPAALKVHGIMALKAYPEALLEIRLIEMP  134
```

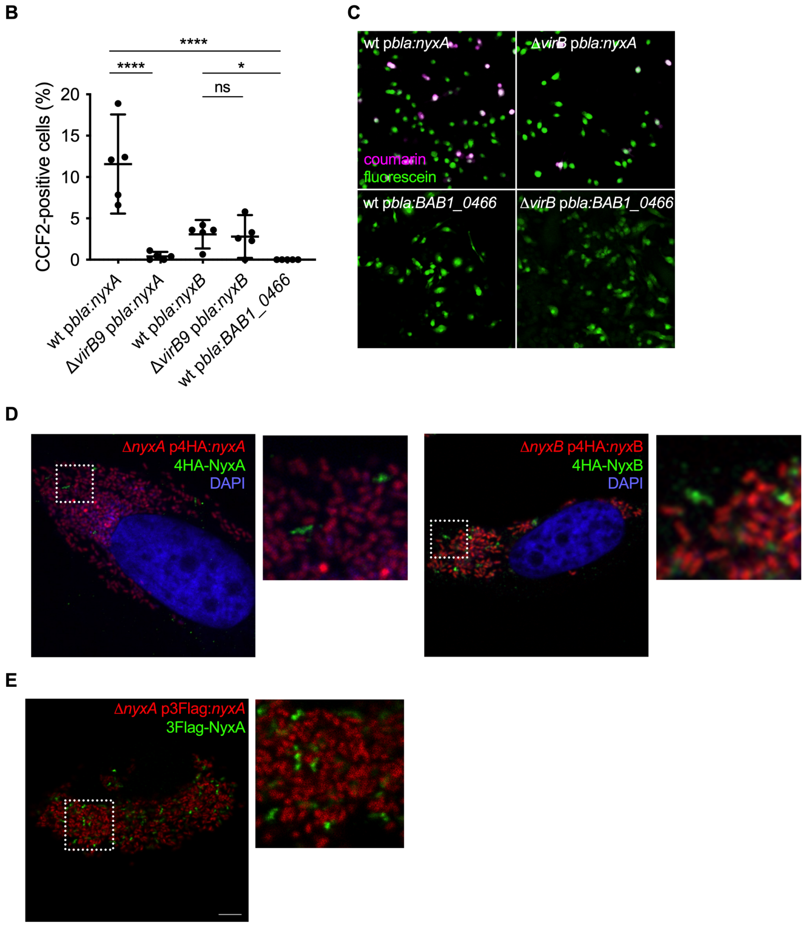


**Fig. S1. NyxA and NyxB are translocated into host cells during infection, accumulating in punctate or filament-like structures. (A)** Amino acid sequence alignment of NyxA and NyxB obtained with ClustalW, matrix EBLOSUM62, showing 82.1% identity, 86.6% similarity and 3.7% gaps. **(B)** RAW macrophage-like cells was infected for 4h with *B. abortus* wild-type or Δ*virB9* expressing TEM1 (encoded by the *bla* gene) fused with NyxA, NyxB or BAB1_0466. The percentage of cells with coumarin emission, which is indicative of translocation, was quantified after incubation with the CCF2-AM substrate. Data represent means ± 95% confidence intervals from 5 independent experiments, with more than 300 cells counted for each condition. **(C)** Representative images for *B. abortus* wild-type or Δ*virB9* carrying p*bla*:*nyxA* or p*bla*:*nyxB* to exemplify the presence of effector translocation visible in coumarin-positive cells (violet) or absence of translocation. **(D)** Representative images to exemplify the filament-like structures visible with both 4HA-tagged NyxA (left) or NyxB (right). HeLa cells were infected for 48h with Δ*nyxA* or Δ*nyxB* strains expressing DSRed and carrying p4HA:*nyxA* or p4HA:*nyxB*. The cell nucleus is visible with DAPI (blue). **(E)** Representative confocal microscopy images showing accumulation of 3Flag-NyxA (green) punctate structures in HeLa cells infected for 48h with Δ*nyxA* expressing DSRed and 3Flag-NyxA.

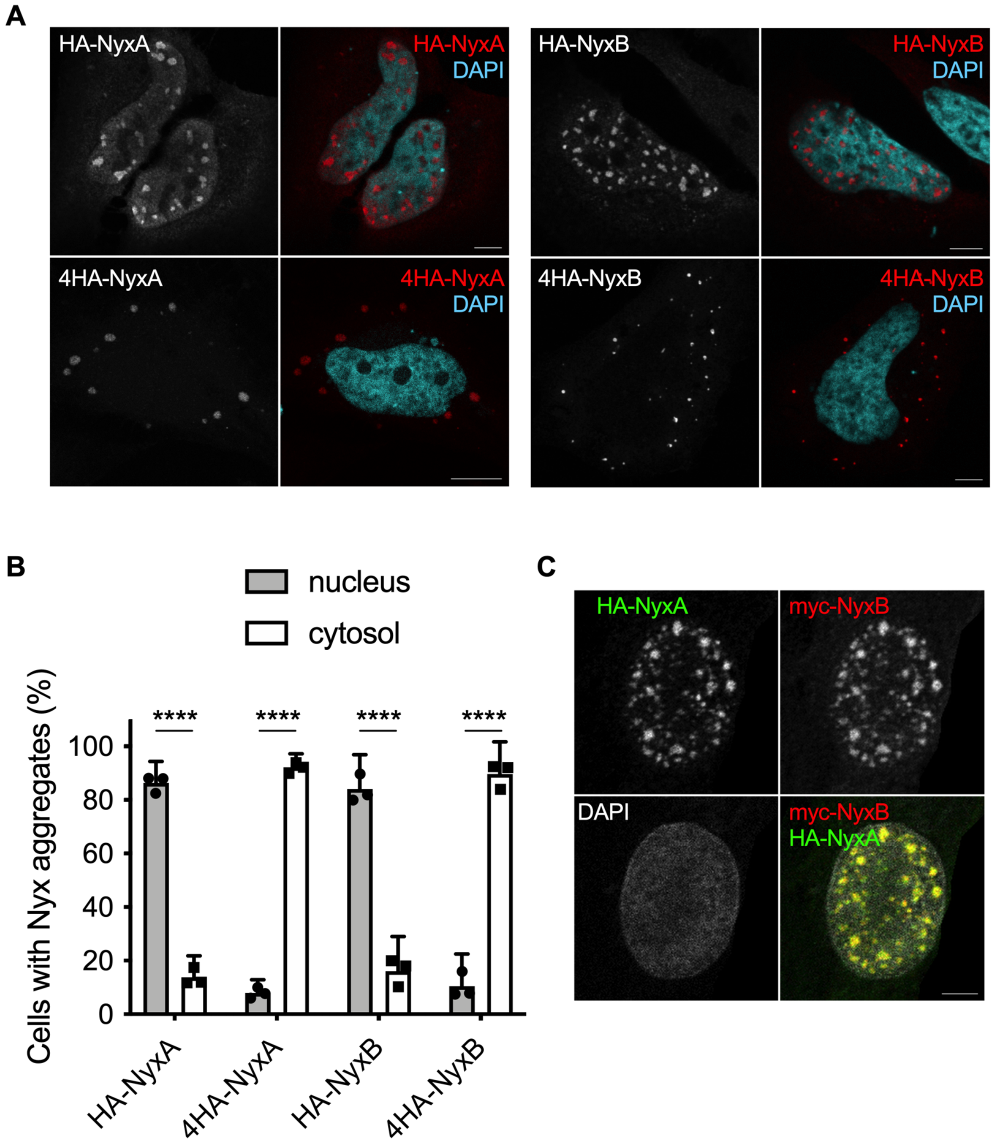


**Fig. S2. NyxA and NyxB target the same cellular compartments. (A)** Representative confocal microscopy images of HA- or 4HA-tagged NyxA and NyxB (red) ectopically expressed in HeLa cells. The nucleus of the cells is labelled with DAPI. Scale bars are 5 µm. **B)** Quantification of the percentage of cells with majority of tagged effector accumulating in cytoplasmic or nuclear structures. Data correspond to means ± 95% confidence intervals from 3 independent experiments. **(C)** Confocal imaging showing co-localisation of HA-NyxA (green) and myc-NyxB (red) aggregates in the nucleus (white).

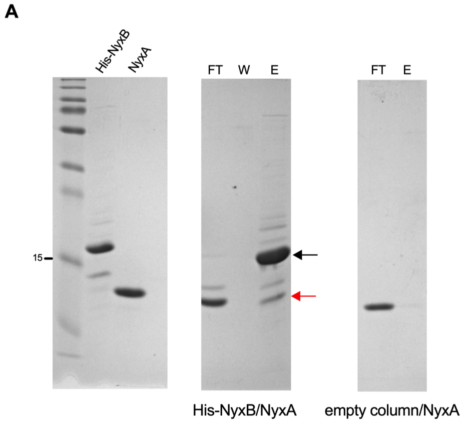


**B**

| Accession | Sum PEP Score | Coverage | # Peptides | # PSMs | # Unique Peptides | # Protein Groups | MW [kDa] | Score Sequest HT |
|---|---|---|---|---|---|---|---|---|
| BnpA | 86,6 | 79,3 | 14 | 99 | 7 | 1 | 15,2 | 150,4 |
| HisBnpB | 26,2 | 38,9 | 9 | 60 | 2 | 1 | 18,8 | 87,0 |

**NyxA coverage:**

GIDPF**TMNAH TNISGSAAFD PNAWHHSQMT IREAIDLSQS GGHPYSSPNV PKGFNTVVGF** FFDCYDWYPA AY**DDEEGNAM**
**KDRELIQYEE W**CAKY**ARKLG LEVKEVEAPA ALKVHGIMTL KAYPEALL**EI RCLEL

**HisNyxB coverage:**

MHHHHHHGKP IPNPLLGLDS TENLYFQGID PFTMNTQATI DTAAVAPLNF DPNAWHHSQM TTLEAIELSR SGGHPY**SSPN**
**VPKGFNTVVG F**FFDTYDWYP AAY**DDEEGNA MKDREL**IQYE DWCAKYARTL **GLEVKEVEAP AALKVHGIMA LKAYPEALLE**
**IRLIEMP**

**Fig. S3. NyxA and NyxB directly interact. (A)** Pulldown experiment using purified NyxA against His-NyxB immobilised on a Ni NTA resin. Empty column was used as a control for non-specific binding. Interactions were visualised with coomassie blue stained gels. The flowthrough (FT), wash (W) and elution (E) fractions are shown for each sample and the molecular weights indicated (kDa). Eluted NyxA and His-NyxB are indicated with black and red arrows, respectively. **(B)** Confirmation of the identity of the two major eluted bands by mass spectrometry. The identified peptides are highlighted in green.

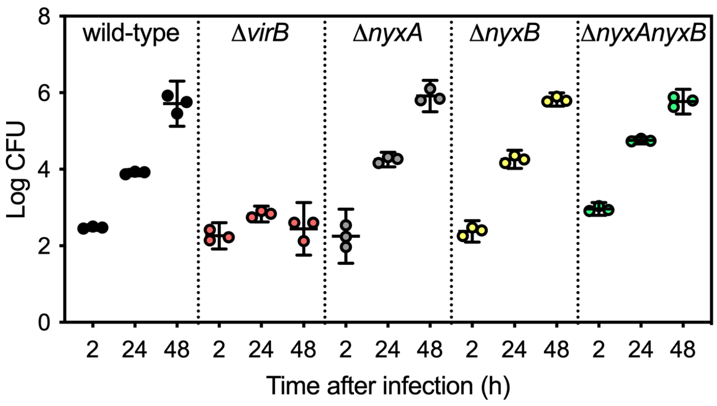


**Fig. S4. Deletion of NyxA and NyxB does not impact intracellular multiplication of *B. abortus*.** Enumeration of bacterial colony forming units (CFU) of wild-type *B. abortus*, Δ*virB9*, Δ*nyxA*, Δ*nyxB* or Δ*nyxAnyxB* following 2, 24 or 48h of infection of HeLa cells.

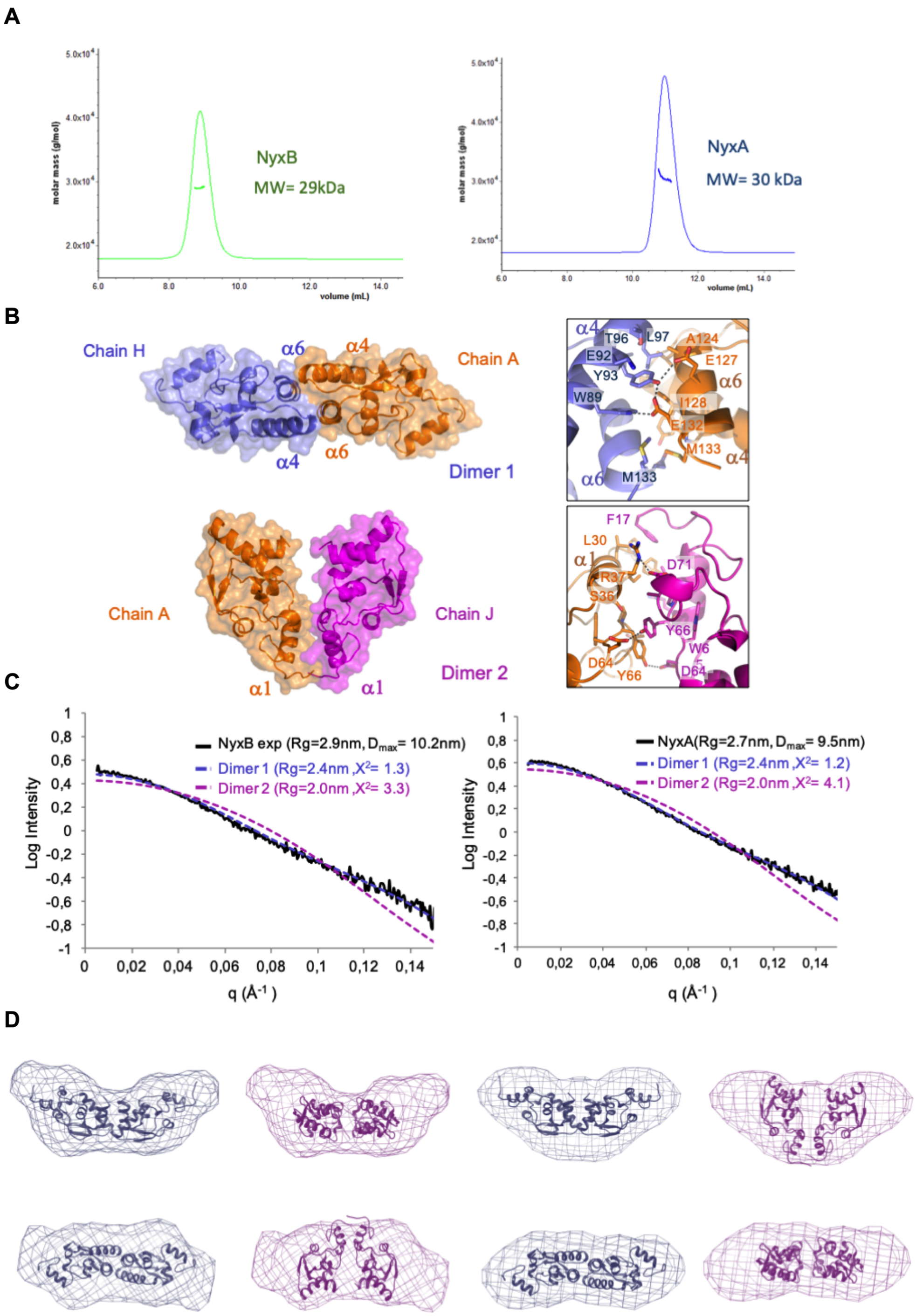


**Fig. S5. NyxA and NyxB dimer formation. (A)** Chromatograms ($A_{280}$, plain) and mass measurements (dots) by Multi Angle Light Scattering of NyxB (left, green) and NyxA (right, blue). The average molecular weight determined is indicated. **(B)** Surface representation of dimer A (chains A and H) and dimer 2 (chains A and J) with detailed view of each interface side chains involved represented as ball-and-sticks. **(C)** Comparison of the theoretical small-angle X-ray scattering profiles of NyxB dimers with experimental data (black curves) obtained for NyxB (left) and NyxA (right). Fitting values (c2) obtained using FoXS server are indicated. **(D)** Fitting of NyxB dimers into *ab initio* SAXS envelopes obtained with GASBOR using NyxB data (left) and NyxA (right) showing that dimer 1 fits much better each of the experimental curves.

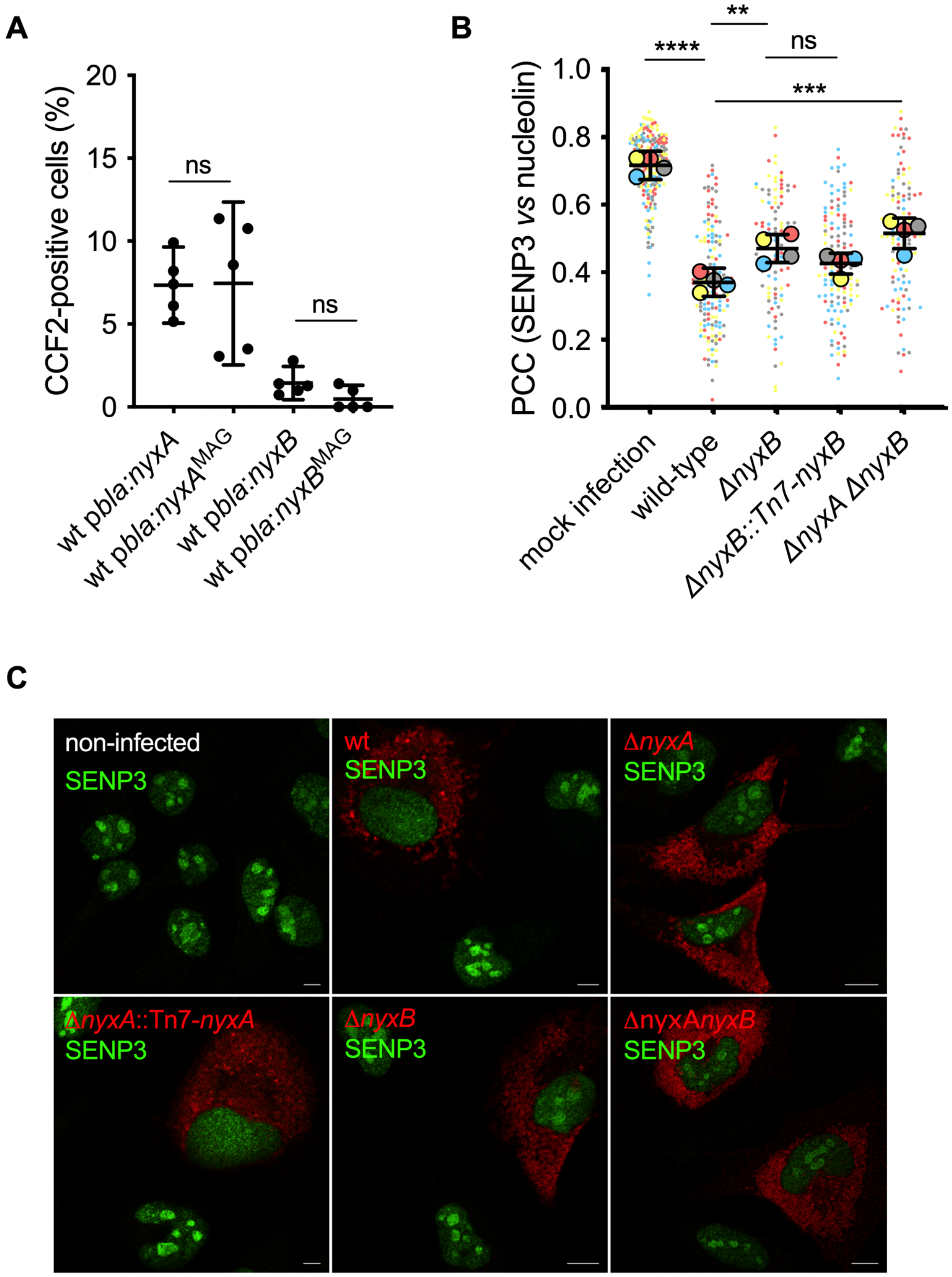


**Fig. S6. The *Brucella* Nyx effectors directly reduce the SENP3 nucleolar localisation in host cells. (A)** RAW macrophage-like cells were infected for 24h with *B. abortus* wild-type expressing TEM1 (encoded by the *bla* gene) fused with NyxA, NyxA$^{MAG}$, NyxB or NyxB$^{MAG}$. The percentage of cells with coumarin emission, which is indicative of translocation, was quantified after incubation with the CCF2-AM substrate. Data represent means ± 95% confidence intervals from 5 independent experiments. **(B)** Quantification of the Pearson's coefficient of SENP3 versus nucleolin in HeLa cells infected for 48h with either *B. abortus* wild-type or Δ*nyxB*, its complemented strain Δ*nyxB*Tn7-*nyxB* or double deletion mutant Δ*nyxAnyxB*. Data are represented as means ± 95% confidence intervals from 4 independent experiments. Each experiment is colour coded and all events counted are shown. Data were analysed using one-way ANOVA by including all comparisons with Tukey's correction. Not all comparisons are shown. **(C)** Representative confocal microscopy images of HeLa cells infected with the different strains expressing DSRed and labelled for SENP3 (green), in comparison to mock infected control cells (non-infected first panel). All scale bars correspond to 5 µm.

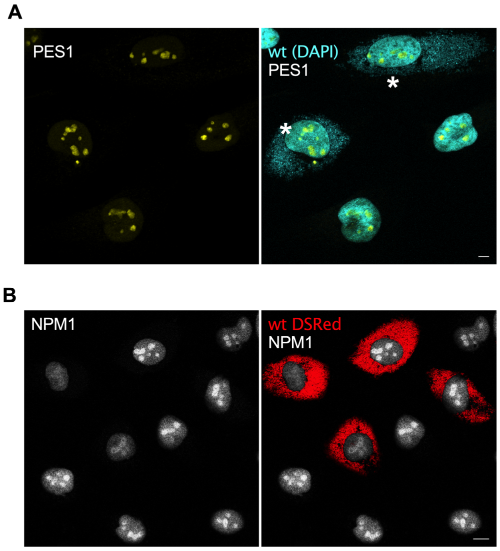


**Fig. S7.** ***B. abortus*** **effect on PES1 and NPM1 nucleolar localization. (A)** Representative confocal images of HeLa cells infected for 48h with wild-type *B. abortus* and labelled for DNA to visualise bacteria and cell nuclei (cyan) and PES1 (yellow). Infected cells are indicated with an asterisk. **(B)** HeLa cells infected for 48h with DSRed-expressing wild-type *B. abortus* and labelled for NPM1 (white). All scale bars correspond to 5 µm.

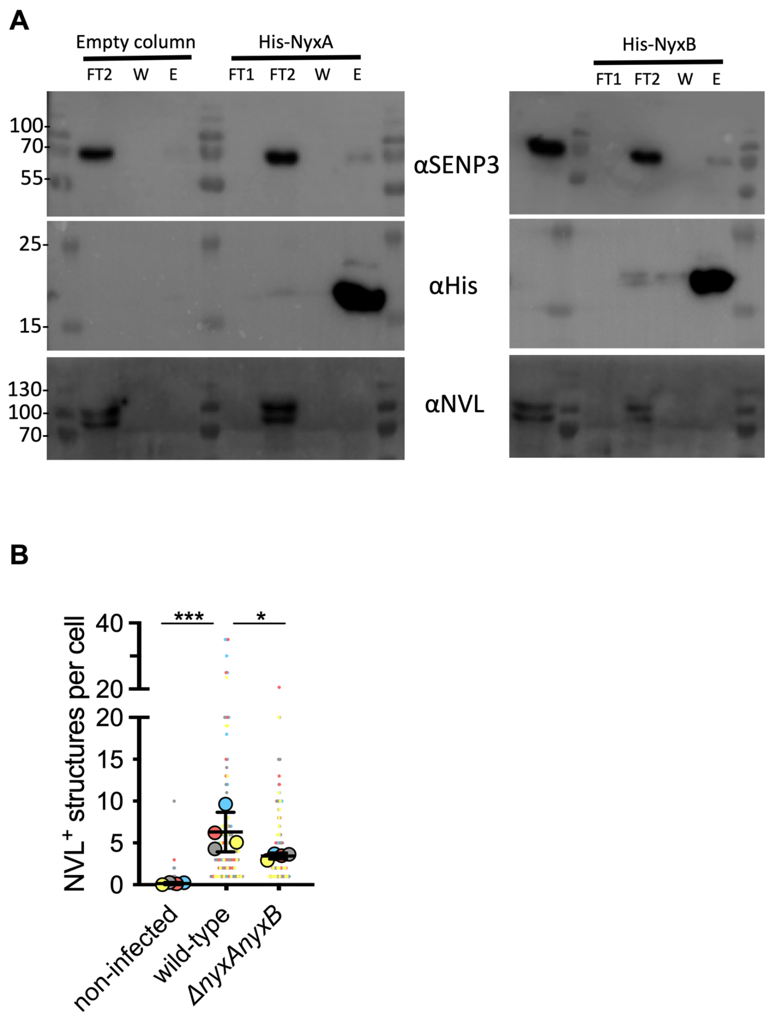


**Fig. S8.** **NyxA and NyxB induce formation of NVL-positive cytoplasmic structures without interacting with NVL. (A)** Pull-down assay with His-NyxA, His-NyxB immobilised on Ni NTA resins that were incubated with a HeLa cell extract. Empty column was used as a control for non-specific binding. Interactions with endogenous SENP3 and NVL were visualised by western blotting using the corresponding antibody, and column binding with anti-His (lower blot). Non-bound fractions (F1 and F2), last wash (W) and elution (E) are shown for each sample and the molecular weights indicated (kDa). The cell extract input is also shown. **(B)** Quantification of the number of NVL-positive cytoplasmic structures in mock infected control iB-MDM in comparison to wild-type or a mutant strain lacking both nyxA and nyxB. Data are represented as means ± 95% confidence intervals from 4 independent experiments. Each experiment is colour coded and all events counted are shown. Data were analysed using one-way ANOVA by including all comparisons with Tukey's correction.

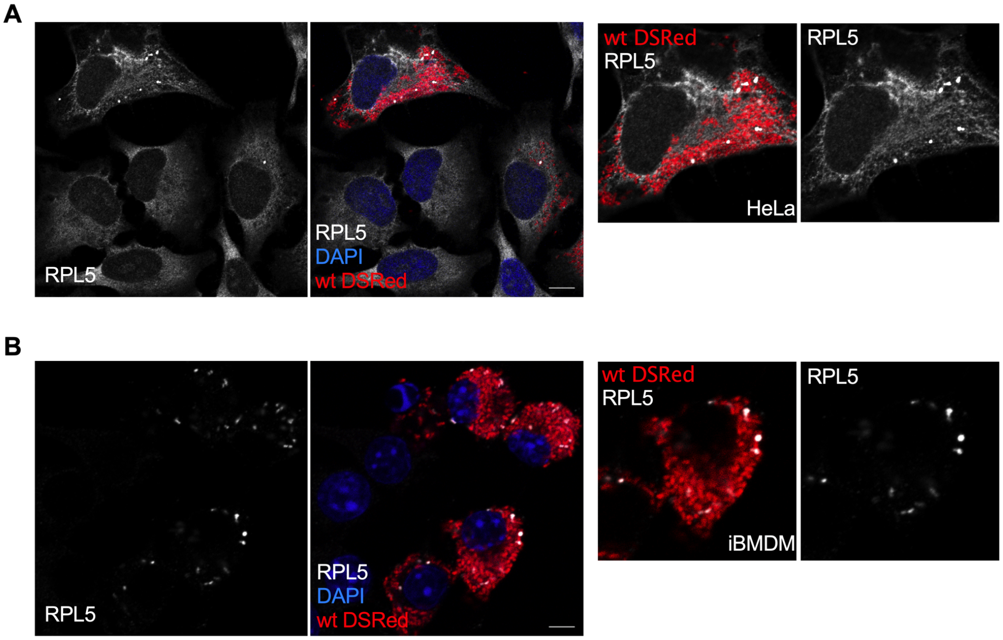


**Fig. S9.** ***B. abortus*** **induces cytoplasmic accumulation of RPL5.** Representative confocal microscopy images of **(A)** HeLa cells and **(B)** iBMDM infected with wild-type DSRed-expressing *B. abortus* for 48h and labelled for RPL5 (white) and DAPI (blue). Zoomed cells are included on the right. All scale bars correspond to 5 µm.

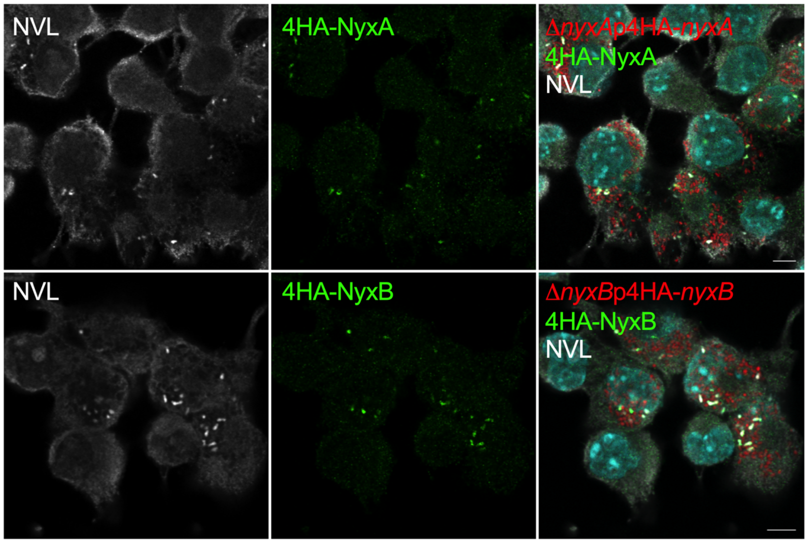


**Fig. S10.** ***B. abortus*** **induces cytoplasmic accumulation of NVL in iBMDM that colocalises with translocated 4HA-tagged NyxA and NyxB.** iBMDM were infected with either *nyxA* expressing DSRed and 4HA-NyxA (top) or *nyxB* expressing DSRed and 4HA-NyxB for 48h and labelled for RPL5 (white) and DAPI (cyan). All scale bars correspond to 5 µm.

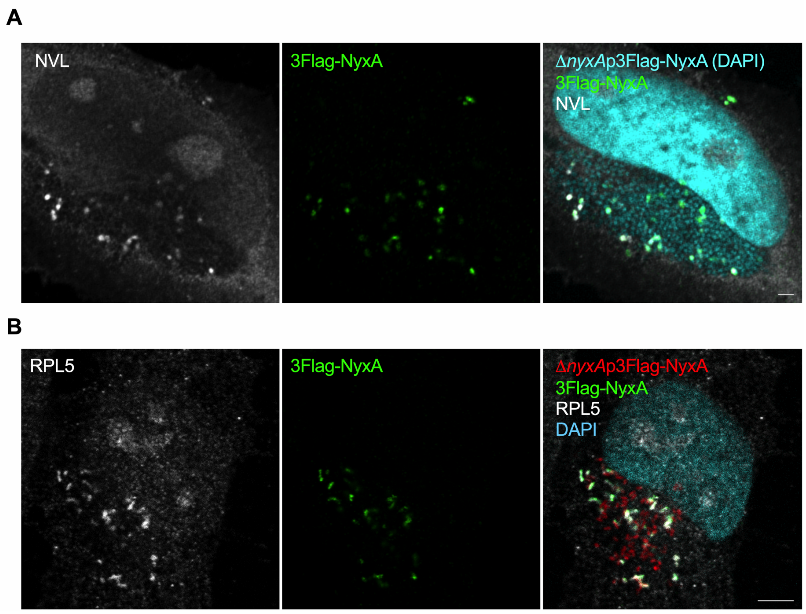


**Fig. S11.** ***B. abortus* induces cytoplasmic accumulation of NVL and RPL5 that colocalises with translocated 3Flag-tagged NyxA.** **(A)** HeLa were infected with *nyxA* expressing 3Flag-NyxA (green) for 48h and labelled for NVL (white) and DAPI (cyan). **(B)** HeLa cells were infected with *nyxA* expressing DSRed and 3Flag-NyxA (green) for 48h and labelled for RPL5 (white) and DAPI (cyan). All scale bars correspond to 5 µm.

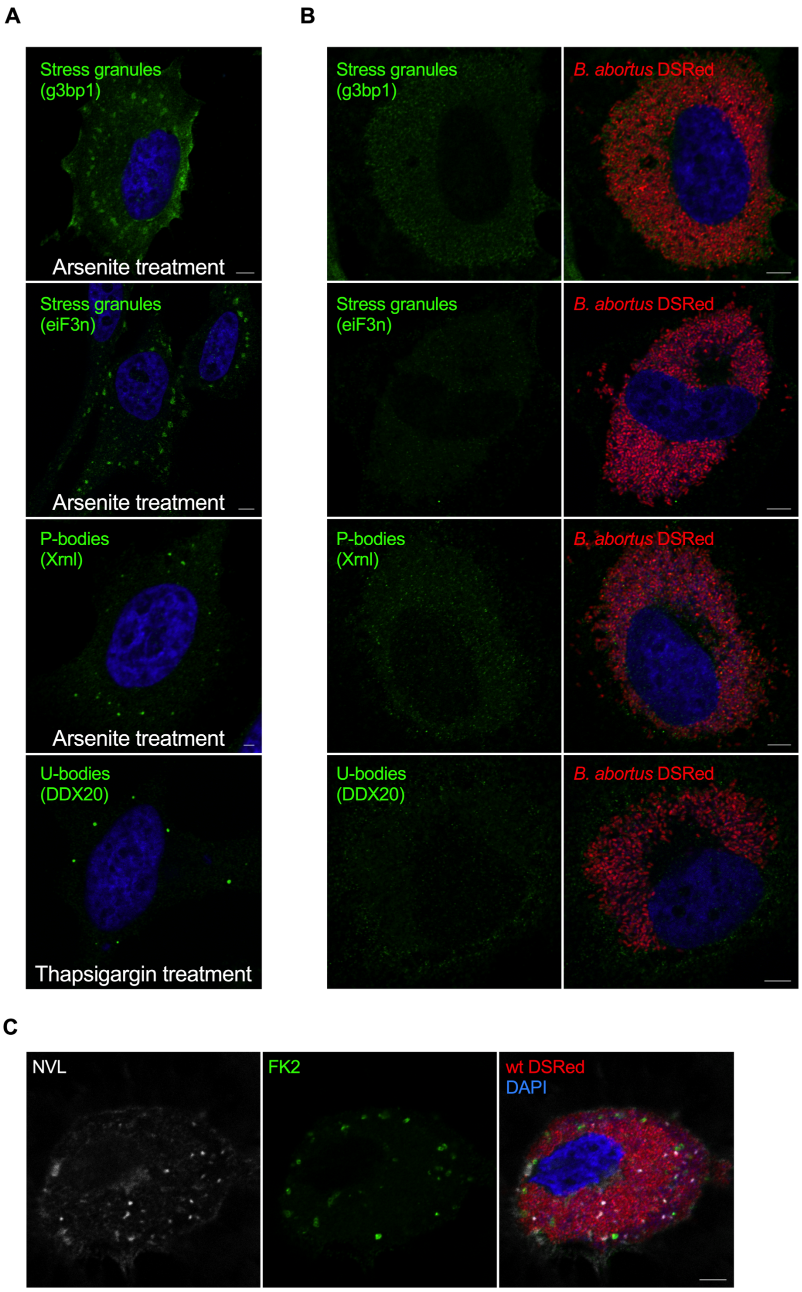


**Fig. S12.** ***B. abortus*** **infection does not induce stress granules, P-bodies, U-bodies. The cytoplasmic NVL structures are not FK2-positive.** HeLa cells were either **(A)** treated with arsenite for 30 min or thapsigargin 4h as positive controls or **(B)** infected for 48h with wild-type DSRed-expressing *B. abortus* and labelled with anti-g3bp1 and eiF3n antibodies to visualise stress granules, anti-Xml antibody for P-bodies and anti-DDX20 antibody for U-bodies. NVL labelling was omitted from the figure for clarity. **(C)** Infected cells were also labelled with the FK2 antibody (green) that recognises mono- and poly-ubiquitinated proteins in addition to NVL (white). All scale bars correspond to 5 µm.

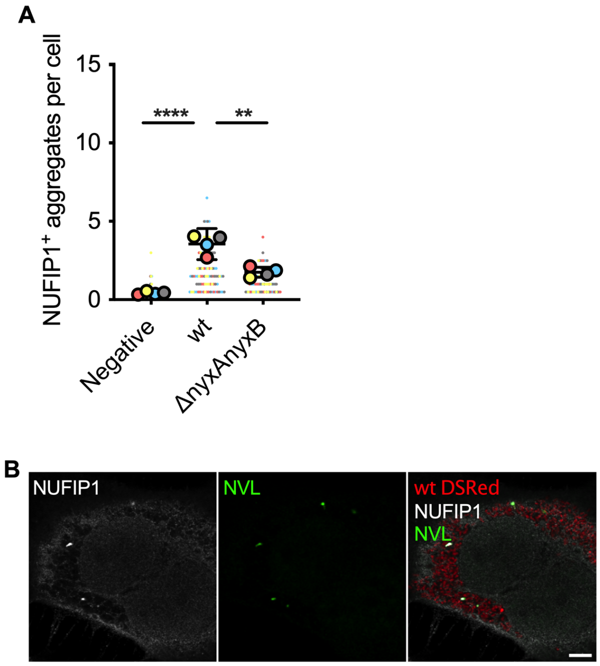


**Fig. S13.** ***B. abortus*** **induces cytosolic shuttling of NUFIP1 in a NyxA/B-dependent manner, notably to Bif. (A)** Quantification of the number of cytoplasmic NUFIP1-positive aggregates at 48h post-infection. Each experiment is colour coded and all events counted are shown. Data were analysed using one-way ANOVA by including all comparisons with Tukey's correction. Not all comparisons are shown. **(B)** Confocal image of HeLa cell infected with wild-type DSRed for 48h and labelled for NUFIP (white) and NVL (green). Scale bar corresponds to 5 µm.

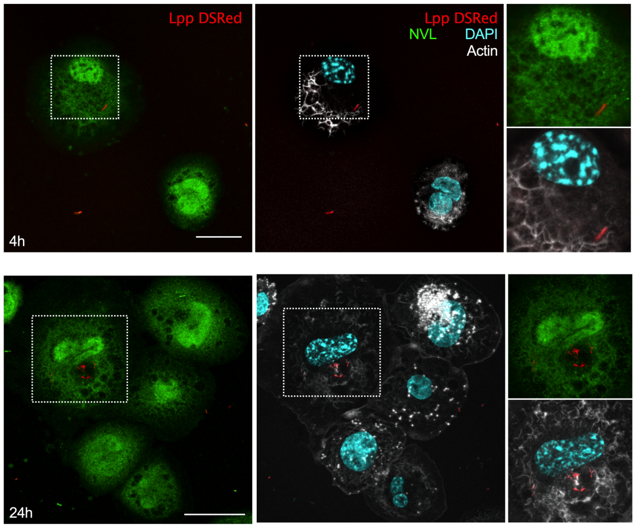


**Fig. S14.** ***Legionella*** **infection does not induce NVL cytoplasmic accumulation.** Immunofluorescence analysis of THP-1 cells infected 4 and 24 hours with wild type *L. pneumophila* strain Paris, carrying a DsRed expressing plasmid. Cells were stained with an anti-NVL antibody (green) and analysed by confocal microscopy. DAPI, light blue; Phalloidin, grey. Scale bars correspond to 10 µm.
